# Dynamics and fate of beneficial mutations under lineage contamination by linked deleterious mutations

**DOI:** 10.1101/068429

**Authors:** Sophie Pénisson, Tanya Singh, Paul Sniegowski, Philip Gerrish

## Abstract

Beneficial mutations drive adaptive evolution, yet their selective advantage does not ensure their fixation. Haldane’s application of single-type branching process theory showed that genetic drift alone could cause the extinction of newly-arising beneficial mutations with high probability. With linkage, deleterious mutations will affect the dynamics of beneficial mutations and might further increase their extinction probability. Here, we model the lineage dynamics of a newly-arising beneficial mutation as a multitype branching process; this approach allows us to account for the combined effects of drift and the stochastic accumulation of linked deleterious mutations, which we call *lineage contamination*. We first study the lineage contamination phenomenon in isolation, deriving extinction times and probabilities of beneficial lineages. We then put the lineage contamination phenomenon into the context of an evolving population by incorporating the effects of background selection. We find that the survival probability of beneficial mutations is simply Haldane’s classical formula multiplied by the correction factor 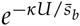, where *U* is deleterious mutation rate, 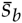 is mean selective advantage of beneficial mutations, *κ* ∈ (1, *ε*], and *ε* = 2 – *e*^−1^. We also find there exists a genomic deleterious mutation rate, 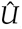, that maximizes the rate of production of surviving beneficial mutations, and that 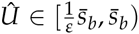. Both of these results, and others, are curiously independent of the fitness effects of deleterious mutations. We derive critical mutation rates above which: 1) lineage contamination alleviates competition among beneficial mutations, and 2) the adaptive substitution process all but shuts down.

---

**B**eneficial mutations are the ultimate source of the genetic variation that fuels evolutionary adaptation, but deleterious mutations are likely to be far more abundant (Muller 1950; Sturtevant 1937). Perhaps for the sake of simplicity, the evolutionary effects of these two types of fitness-affecting mutations were generally considered separately in early studies. For example, Muller (1964) assumed that beneficial mutations were negligible and reasoned verbally that deleterious mutations should have disastrous consequences for populations in the absence of recombination because of the recurrent, stochastic loss of genotypic classes with the fewest deleterious mutations–Muller’s ratchet (Felsenstein 1974). Haldane (1927), on the other hand, focused on the fate of single beneficial mutations in the absence of other fitness-affecting mutations and used single-type branching process theory to show that most such beneficial mutations are lost to what is now called genetic drift: the fixation probability of such a beneficial mutation is only about twice its selective effect, *s_b_*, for small *s_b_*.

In reality, of course, multiple fitness-affecting mutations (both beneficial and deleterious) can be present simultaneously in populations, and these mutations can influence each others’ fates and evolutionary effects as a consequence of linkage (reviewed in Gordo and Charlesworth (2001); Charlesworth (2013, 2009); Barton (2009)). Interactions between beneficial and deleterious mutations are of particular interest in this regard, because such interactions-in contrast to interactions between beneficial mutations alone–can determine whether a population will increase or decrease in fitness. Indeed, recent studies (Bachtrog and Gordo 2004a; Poon and Otto 2000; Goyal *et al*. 2012; Kaiser and Charlesworth 2009; Silander et *al*. 2007) have indicated that beneficial mutations (including reversions of deleterious mutations) can impede or halt the fitness loss predicted in asexual populations under Muller’s ratchet, as originally suggested by Haigh (1978a). Moreover, a number of theoretical studies (Johnson and Barton 2002a; Jiang et *al*. 2011; McFarland et *al*. 2014; Bachtrog and Gordo 2004a; Charlesworth 2013; Peck 1994) have shown that Haldane’s classical fixation probability of 2*s_b_* for a beneficial mutation can be reduced by the effects of selection against linked deleterious mutations (Good and Desai 2014; Hartfield *et al*. 2010; C W Birky and Walsh 1988): in principle, such effects include both background selection against deleterious mutations already present in the genome on which the beneficial mutation appears, and selection against deleterious mutations that arise and accumulate in genomes carrying the beneficial mutation. The latter form of selective effect has not previously been analyzed in isolation; it is the primary focus of the current paper and will be referred to as *lineage contamination*.

In preliminary computer simulations, we observed that the fixation probability of a beneficial mutation appearing in an otherwise initially homogeneous asexual population with a high genomic mutation rate is considerably reduced below Haldane’s classical 2*s_b_* expectation. We hypothesized that the lower probability of fixation of a beneficial mutation in this situation can be attributed to lineage contamination: specifically, Muller’s ratchet operates at a much faster rate in the small lineage founded by the beneficial mutation than in the rest of the population. Here, we present the results of analytical modeling and further computer simulations that support this hypothesis and show how lineage contamination affects fixation probabilities, dynamics of differential load (relative fitness dynamics), sojourn dynamics, and fitness effects of surviving beneficial mutations.

We model the influences of background selection and lineage contamination, both singly and jointly, on the fate of beneficial mutations. Under background selection alone, a beneficial mutation that lands on the best genetic background (the one least loaded with deleterious mutations) always has a non-zero probability of achieving fixation in an asexual population. In contrast, under lineage contamination alone, a beneficial mutation can have a probability of fixation that is zero if the mutation rate is high enough. Our simulations and analytical results suggest that when both background selection and lineage contamination are operating – as they do in real populations – asexual populations traverse a continuum of evolutionary regimes as the genomic mutation rate increases: at low mutation rates, beneficial mutations appear infrequently enough that they do not interfere with each others’ progress to fixation (the “periodic selection” regime; Sniegowski and Gerrish (2010)); as the mutation rate increases, alternative beneficial mutations begin to compete with each other (the “clonal interference” regime); as mutation rate increases further, we find that lineage contamination can suppress a fraction of beneficial mutations that is sufficient to cause a population to revert to the periodic selection regime; ultimately, at very high mutation rates, a regime can be reached in which beneficial mutations are no longer substituting. Significantly, these last two regimes would not obtain without the operation of lineage contamination: at high mutation rates, background selection alone cannot shut down clonal interference or the adaptive substitution process, but lineage contamination can. Our results, therefore, indicate that the lineage contamination effect is central to determining the adaptive fate of a population when both beneficial and deleterious mutations are arising (Bull and Wilke 2008; Bull *et al*. 2007; Springman *et al*. 2009).

## Beneficial lineages in a homogeneous population

We model the random accumulation of deleterious mutations within a growing lineage founded by the occurrence of a beneficial mutation (henceforth, *beneficial lineage*). Our main objective in this section is to study the effects of lineage contamination in isolation, and to this end we model beneficial lineages arising within initially homogeneous populations.

We are interested in how the accumulation of *linked* deleterious mutations affects the dynamics and fate of beneficial mutations; as a first approximation, we assume complete asexuality. We assume that relative fitness effects of mutations do not change over the relevant time span, i.e., the environment remains constant over this time span, and there are no frequencydependent effects other than the one examined here (due to differential rates of Muller’s ratchet). All models assume that fitness effects of mutations are multiplicative, i.e., no epistasis. Finally, our multitype branching process model assumes that reproduction is by binary fission (e.g., bacteria, cell lines): thus, individuals can have a maximum of two offspring. Simulations that relax this assumption give qualitatively similar results; quantitatively, however, fixation probabilities derived from the binary fission model may be approximately halved for the more general model in which numbers of offspring are Poisson-distributed.

Finally, it will facilitate further reading to precisely define three terms: *extinction probability*, 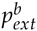, is the probability that a beneficial lineage, arising in an otherwise infinite population, becomes extinct in finite time (the superscript *b* indicates that this probability pertains to the beneficial lineage in question and not the whole population); *survival probability* is the complement of the extinction probability: 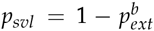; finally, *fixation probability, p_fix_*, is the probability that a lineage will displace the rest of a finite population (becomes fixed) in finite time. We note that we have dropped the superscript in *p_svl_* and *p_fix_*, as reference to the beneficial lineage is implied. We further note that it is possible for an ultimately doomed lineage to become fixed in a finite population, implying *p_fix_* > *p_svl_*.

### Multitype branching process model

Our stochastic model is a discrete-time multitype branching process, where a “type” 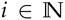 corresponds to the number of acquired deleterious mutations. The model describes the evolution of the composition of the population **X**_*t*_ = (*X*_*t*,0_, *X*_*t*,1_,…), *X_t,i_* being the number of individuals carrying *i* deleterious mutations at time 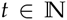. We denote by *U* ⩾ 0 the deleterious mutation rate and by 0 ⩽ *s_b_* ⩽ 1 and 0 < *s_d_* < 1 the selective advantage of beneficial mutations and disadvantage of deleterious mutations, respectively. The model can be described as follows: at each time-step, each individual produces two descendants carrying as many deleterious mutations as itself. Each descendant might accumulate during this reproduction *k* additional deleterious mutations, with probability *e^−U^U^k^*/*k*!. If the parent was of type *i*, the descendant is then of type *i* + *k* and is selected according to its fitness, i.e. with probability proportional to (1 + *s_b_*)(1 − *s_d_*)^*i*+*k*^. Therefore, an individual of type *i* produces a total number of 0, 1 or 2 descendants, each of them being of a type greater than or equal to *i*. We refer to the Supporting Information (SI) for a more detailed description of the model.

We consider a sub-population carrying a beneficial mutation (i.e., a single beneficial lineage) arising in a large wild-type population. In order to study the lineage contamination effect in isolation, we assume that both populations initially do not carry any deleterious mutations. For this purpose we consider two independent branching processes: 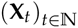 describing the evolution of a wild-type population of initial size *N*, hence with *s_b_* = 0 and initial state **X**_0_ = (*N*,0,0,…), and 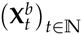 describing the evolution of a single beneficial lineage, with *s_b_* > 0 and initial state 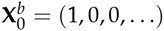.

### Mean demographic dynamics of each sub-population

The mean wildtype population size (all types combined) at time *t* is given by *Ne*^−Ut^*e*^*U*(1−(1−*s_d_*)^*t*^)(1−*s_d_*)/*s_d_*^ (S4). A beneficial mutation occurring within the wildtype population founds a beneficial lineage whose mean size is given by (1 + *s_b_*)^*t*^ *e^−Ut^e*^*U*(1 −(1−*s_d_*)^*t*^)(1−*s_d_*)/*s_d_*^ (S5). Note that as time tends to infinity this quantity tends to +∞ if *U* < ln (1 + *s_b_*), to *e*^*U*(1−*s_d_*)/*s_d_*^ if *U* = ln (1 + *s_b_*), and to 0 if *U* > ln (1 + *s_b_*). The latter convergence will typically not be monotonic (Figure S1).

### Extinction and survival probabilities

The previous result concerning the evolution of the mean beneficial lineage population size can be refined by looking at the *extinction prohahility* of the beneficial lineage. By this we mean the probability 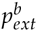 that the process 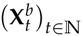 does become extinct, i.e. 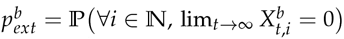. We show (Proposition 1; see SI) that although the number of types is infinite, this probability also equals 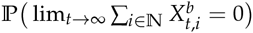. The beneficial lineage almost surely becomes extinct if and only if *U* ⩾ *U_c_*, where the critical deleterious mutation rate is given by

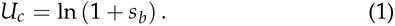

Of course, this implies that if *U* < *U_c_* then 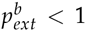, i.e. the beneficial lineage can survive with positive probability. We find that survival probability 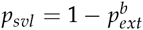 is bounded by:

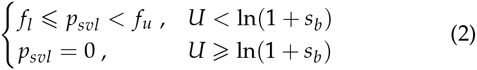

where

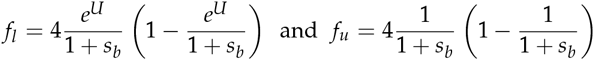

Lower bound *f_l_* is achieved when *s_d_* > *s_b_*, and the upper bound *f_u_* is achieved when *s_d_* → 0. Exact computation of *p_svl_* is achieved numerically using algorithm (S11) derived in the SI. Figure 1 plots examples of such computations (thin intermediate curves) as well as limiting cases *f_l_* and *f_u_* (thick curves) as a function of the deleterious mutation rate *U* and selective advantage *s_b_*, respectively.

**Figure 1.**
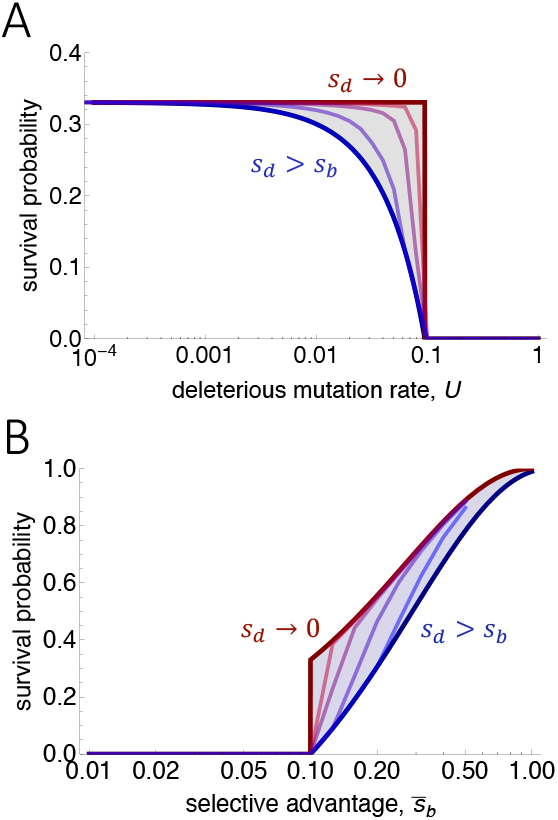
Survival probabilities as functions of: (A) deleterious mutation rate, and (B) selective advantage of the beneficial mutation. Solid curves plot survival probabilities of a beneficial mutation in an otherwise homogeneous population (lineage contamination in isolation). Blue curves plot the case *s_d_* > *s_b_*, red curves plot the limit *s_d_* → 0, and curves of intermediate color plot survival probabilities for intermediate values of *s_d_* (numerical solutions to the survival probability equations given in the SI). Parameters are: *s_b_* = 0.1 (for A), and *U* = 0.1 (for B).

### Fixation probabilities

In this branching process model the *fixation prohahility* of the beneficial lineage is the probability *p_fix_* that at some point the whole population carries the beneficial mutation. Note that because we take into account stochastic variation in population size, the beneficial mutation might not be *permanently* established even after fixation, because the population might eventually become extinct afterward. The fixation probability corresponds here exactly to the probability that the wild-type population dies out before the single beneficial lineage does: 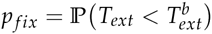, where 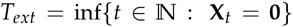 is the extinction time of the wild-type population (alternatively, beneficial lineage). From what precedes we know that *T_ext_* is almost surely finite, whereas 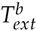 is almost surely finite if and only if *U* ≾ *U_c_*. Note also that because of the strict inclusion of the probability events 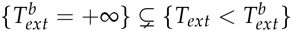, we know that 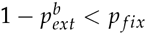. This implies in particular that in this model the fixation probability is never zero. Although we cannot provide a closed-form expression for *p_fix_*, this probability can be well approximated numerically by Equations (S12) - (S13). We illustrate this result in Figure S5 where we plot *p_fix_* as a function of the deleterious mutation rate.

### Fitness dynamics of a beneficial lineage within a population

The fitness of the beneficial lineage at time *t* is given by

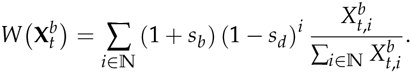

Because of the potential extinction of the population, the random variable 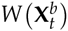 is only defined for 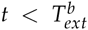 similarly define the fitness of the whole population 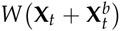, and focus our study on the dynamics of the relative fitness 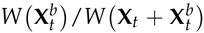. Because we assume in our model that the wild-type population is initially large, we approximate the relative fitness by its almost certain limit as *N* tends to infinity (S14), namely

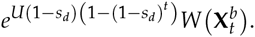

We also prove that the mean value of this relative fitness tends as time tends to infinity to (1 + *s_b_*) (1 − *p_ex_t*) (S15)-(S16). A plot of this long-term limit is given in Figure S6. In order to have a more accurate description of the evolution over time of the relative fitness, we provide in addition an upper and lower bound (S17)-(S18) of its mean value for each 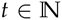, as illustrated in Figure 2.

**Figure 2.**
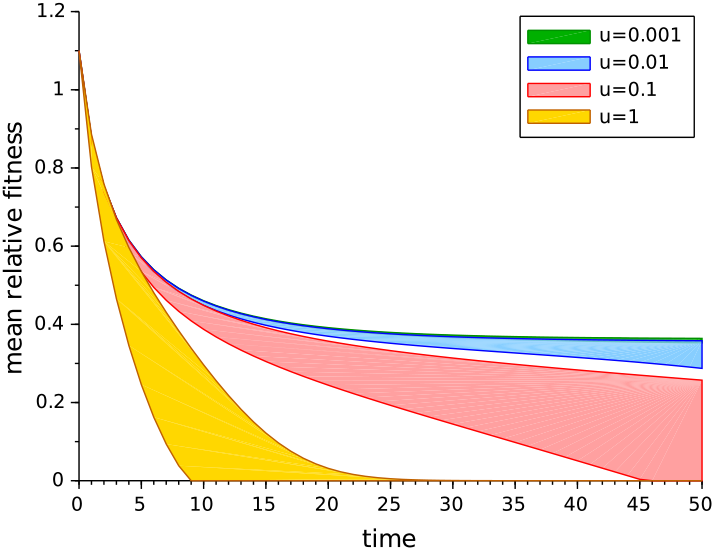
Evolution over time of the mean relative fitness of a single beneficial lineage (*s_b_* = 0.1) in a large wild-type population, initially homogeneous, with *s_d_* = 0.03. Upper and lower bounds were obtained as described in the SI. The decrease in relative fitness is caused by the comparatively rapid accumulation of deleterious mutations owing to the small size of emerging beneficial lineages, i.e., lineage contamination.

### Mutational meltdown of a beneficial lineage

Our goal here is to study the synergy between the loss of the least-loaded classes and the potentially decreasing size of the beneficial lineage. For technical reasons detailed in the SI, we consider in this section the continuous-time analog 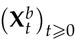 of the branching processes 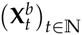 studied previously. Assuming that at time *t* the least-loaded class in the beneficial lineage’s population is of type *i*, the process at this time is of the form 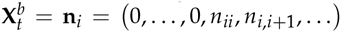. Conditionally on 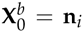, we define the extinction time of the least-loaded class as 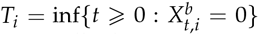. The mutational meltdown effect is then fully described by the sequence of random variables *T*_0_, *T*_1_,…

Note that *T*_1_ strongly depends on the random value 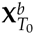 taken by the process at the beginning of the time interval [*T*_0_, *T*_0_ + *T*_1_]. Note also that assuming 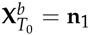, the strong Markov property enables study of the process on the latter interval to be reduced to its study on [0, *T*_1_], conditionally on 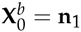. We thus provide in Proposition 2 (SI) an explicit computation of the cumulative distribution function 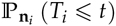 of the time to extinction of the least-loaded class of type *i*, for any *i* and any initial condition 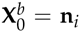. From this we deduce its mean value 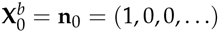. Again, three different regimes appear depending on whether *U* < *U_c_*, *U* = *U_c_* or *U* > *U_c_*. We illustrate this result in Figures S2 and S3, where we plot the cumulative distribution function and mean value of the extinction time *T*_0_ of the first least-loaded class, with 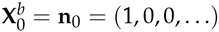.

Finally, in order to study not only the behavior of each extinction time separately but to take into account the stochastic evolution of the process 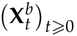, we compute the sequence of the mean extinction times 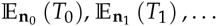, where the deterministic sequence 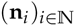 is chosen to reflect as accurately as possible the mean evolution of 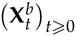. We naturally choose **n**_0_ = (1,0,0,…), and then define **n**_1_ as the mean value of the process at the end of the first time interval [0, *T*_0_]. Because this mean value might not be integer-valued, we round each of its coordinates to the closest integer. Hence we set 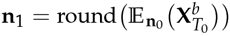, and iteratively define in a similar manner **n**_2_, **n**_3_,…. As proved in Proposition 2 (SI), we can explicitly compute each 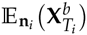, which combined with the previously mentioned computation of 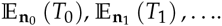 for any initial condition **n**_*i*_, enables us to obtain the desired sequence 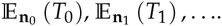. Figure 3 illustrates this result and provides a visualization of the mutational meltdown effect in a single beneficial lineage for different values of *U*, *s_b_* and *s_d_*.

**Figure 3.**
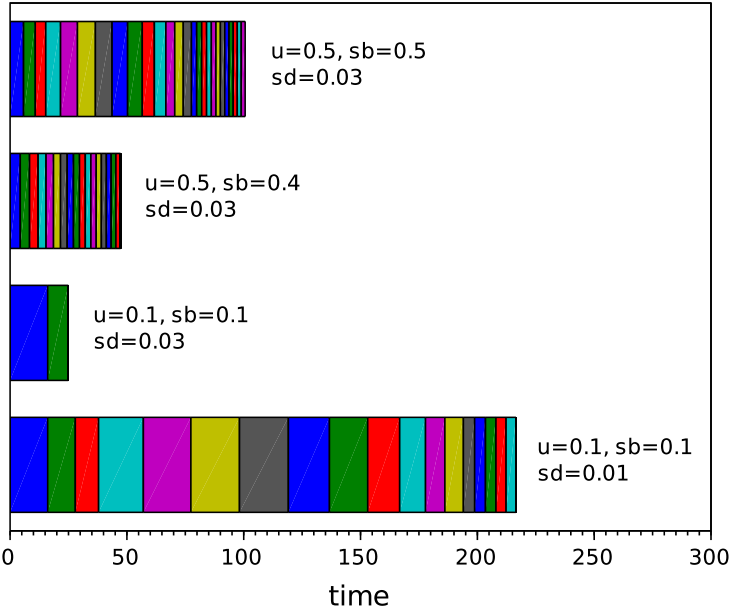
Mutational meltdown of beneficial lineages. Sequence 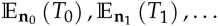 of the mean extinction times of the least-loaded classes, for one single beneficial lineage.

## Beneficial lineages in an evolving population

Until now, we have examined the process of lineage contamination in isolation; that is, the accumulation of deleterious mutations occurring *after* the production of a beneficial mutation. In addition, we have assumed that we know the selective advantage of the focal mutation.

In real populations, however: 1) deleterious mutations can occur both after and before the appearance of a beneficial mutation, and 2) we generally will not know the selective advantages of beneficial mutations. Deleterious mutations that appear before create a deleterious background upon which the beneficial mutation arises; selection against this deleterious background is *background selection* (Charlesworth *et al*. 1993; Stephan 2010). Here, we model the growth and fate of beneficial mutations of varying selective advantages arising in a population already contaminated with deleterious mutations.

Angled-bracket notation in this section indicates average over all possible trajectories, or “states”, (*ensemble* average) of a beneficial lineage emerging in an otherwise heterogeneous (evolving) population. (The absence of angled-brackets indicates that the focal beneficial lineage arises in an otherwise homogeneous population, as in the previous section.)

### Incorporating background selection

If a beneficial mutation is produced on a background carrying *j* deleterious mutations, the initial growth rate of the resulting lineage is:

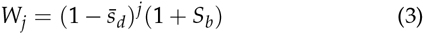

where *S_b_* is a random variable denoting the selective advantage of the beneficial mutation, with mean 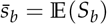. Essentially, to incorporate background selection, we simply replace 1 + *s_b_* in the previous section with *W_j_*.

Of course, we do not know beforehand how many deleterious mutations will be present in the background upon which a beneficial mutation arises. But we do have accurate predictions for both the average number of deleterious mutations in the population, as well as the probability that a beneficial mutation will arise on a background with a given number of deleterious mutations.

When mutation rates are low and population sizes are large, classical theory (Haigh 1978a; Johnson 1999b) predicts that individuals in a population will acquire a Poisson-distributed number of deleterious mutations with parameter 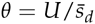. For our purposes, the assumptions of low mutation rate and large population size may be too restrictive, as we wish to explore effects of high mutation rates in finite populations. For exact computation of results, therefore, we will rely on the more encompassing results derived by Gessler (1995) that relax these assumptions, giving the probability of a background having *j* mutations (re-derived in SI):

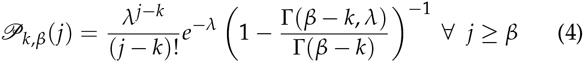

where *λ* = *θ* − *k*, and *k* and *β* are integers defined by 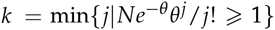, and 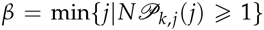. To derive approximate analytical expressions, where defensible, we nevertheless resort to the straight Poisson distribution from classical theory, the rationale being that tail probabilities lower than 1/N will have negligible effects on the quantities being derived.

### Survival probability

The ensemble-averaged probability of survival is bounded by:

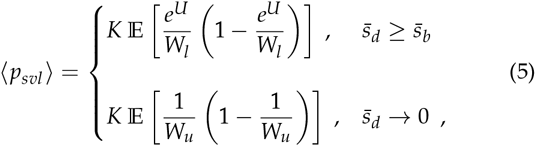

where *K* = 4 for binary fission, and *K* ≈ 2 for Poisson-distributed offspring; *W_l_* = (1 − *s_d_*)^*Jl*^ (1 + *S_b_*) and 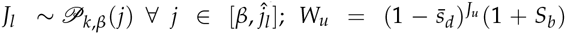 and 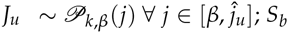 is also a random variable and, for each value of 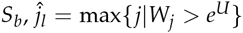 and 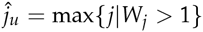, and *W_j_* is defined by Equation (3). Figure 4 plots exact calculations of *p_svl_* and of 〈*p_svl_*〉 by fixing *S_b_* = *s_b_*. Monte Carlo integration of (5), in which *S_b_* was drawn from an exponential distribution with mean 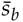, gives probabilities that are indistinguishable from the approximations we now derive.

**Figure 4.**
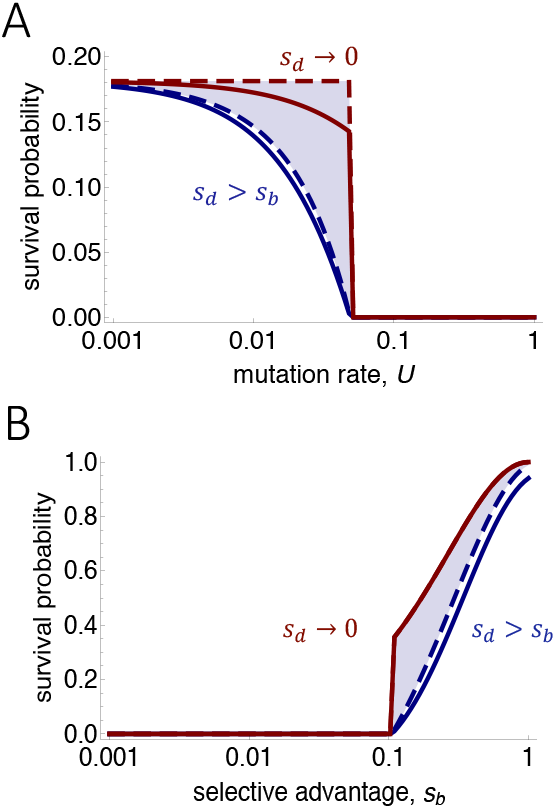
Survival probabilities, computed exactly from (5) by fixing *S_b_* = *s*_b_, plotted as functions of: (A) deleterious mutation rate, and (B) and selective advantage of the focal beneficial mutation. Solid curves plot survival probabilities of a beneficial mutation in a heterogeneous (evolving) population. Dashed curves plot survival probabilities in an otherwise homogeneous population (lineage contamination in isolation). Blue curves plot the case 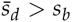, and red curves plot the limit 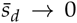. Parameters are: *N* = 50000, *s_b_* = 0.05 (for A), and *U* = 0.1 (for B).

### Approximate survival probability

This approximation is suggested by the observation (Figure 4) that fixation probability increases sharply at the critical selective advantage, 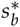, above which fixation probability becomes positive. We thus explored the possibility that fixation probability might be approximated as simply the probability that 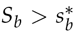 times the probability that the beneficial mutation survives. If the beneficial mutation in question arises on a background with *j* deleterious mutations, then 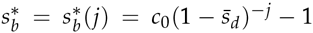, where *c*_0_ = 1 for 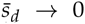 and *c*_0_ = *e^U^* for 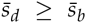, and 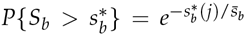. Taking the logarithm of this probability, multiplying by the corresponding Poisson probabilities and summing over *j*, we employ Jensen’s inequality to derive expressions providing exact minimums on both upper and lower bounds for the ensemble survival probability:

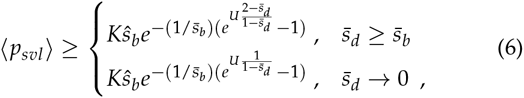

where 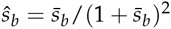. This expression is a bound-of-bounds and thus of questionable utility. Comparison with simulations reveals the upper bound (for 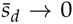) to be quite accurate but the lower bound (for 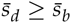) to be overly conservative.

Employing a different approach that does not rely on Jensen’s inequality (SI), we find that, to a very good approximation:

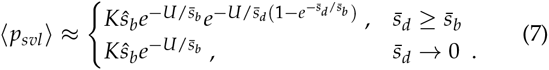

From this, it is apparent that the smallest value of 〉*p*_svl_〈 is achieved when 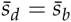, so that survival probability is bounded as:

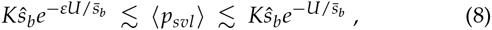

where *ε* = 2 − *e*^−1^ ≈ 1.63. Remarkably, the foregoing bounds on survival probability are independent of 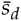. Comparison with simulations (Figure 5) reveals that the bounds given by Equations (7) and (8) are very accurate. From (7), we can see that the upper bound in (8) is approximated under the wider range of circumstances, because: 1) when 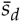 is small the upper bound is approximated, and 2) when 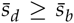 the lower bound obtains, but as 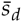 becomes increasingly larger than 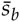, 〉*p_svl_*〈) moves away from the lower bound and back towards the upper bound. There is nevertheless a restricted range of values for 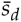 – namely when 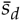 is close to 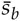 – over which the lower bound obtains.

**Figure 5.**
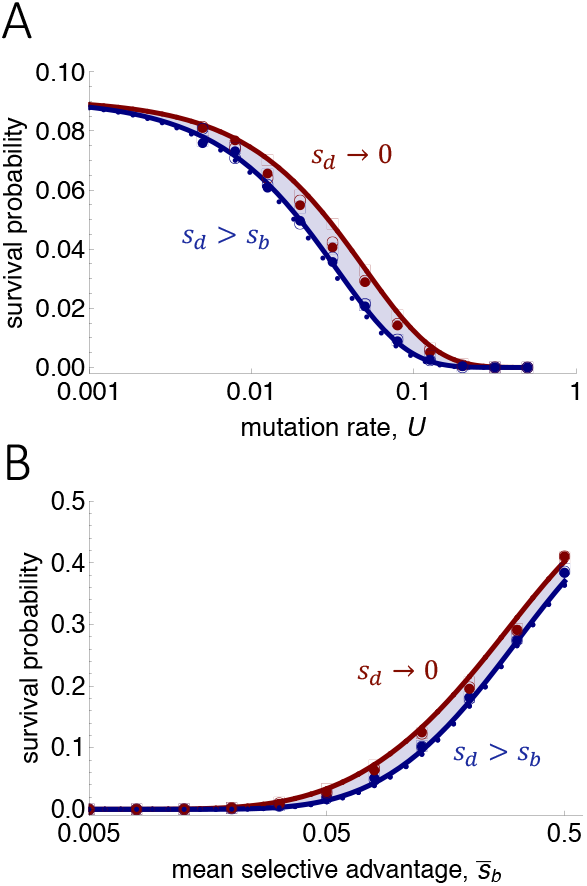
Ensemble survival probabilities. Solid curves plot Equation (7); dotted curves plot Equation (8). Red dotted curves are not visible because they are indistinguishable from solid red curves; blue dotted curves are very close to solid blue curves. Points each plot the fraction of 15,000 individual-based stochastic simulations in which the focal beneficial mutation survived. Lineage survival was defined as either achieving a size of 5000 or lasting for 5000 generations. Solid points plot simulation results for which no further beneficial mutation occurred (*c* = 0); open circles plot simulations that allowed further beneficial mutations to occur at rate *c* = 0.001 times the deleterious rate *U*; open squares plot simulations for which *c* = 0.01. For panel A, beneficial mutations have mean selective advantage 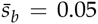; for panel B deleterious mutation rate is *U* = 0.05. When 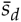 is small, the upper bound obtains (red curve); when 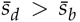, the lower bound obtains (blue curve). In simulations, red points plot the case 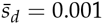 and blue points plot the case 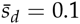.

### Selective advantages of surviving beneficial mutations

Because of lineage contamination, beneficial mutations of small effect will have a very small or zero chance of survival; beneficial mutations that do survive, therefore, will tend to be of larger selective advantage. Following logic similar to that of the previous subsection, we derive the ensemble survival probability for a beneficial mutation of given selective advantage *s_b_*. Given this survival probability, expected selective advantages of surviving beneficial mutation rates, as well as approximate bounds, are plotted in Figure 6 as a function of deleterious mutation rate.

**Figure 6.**
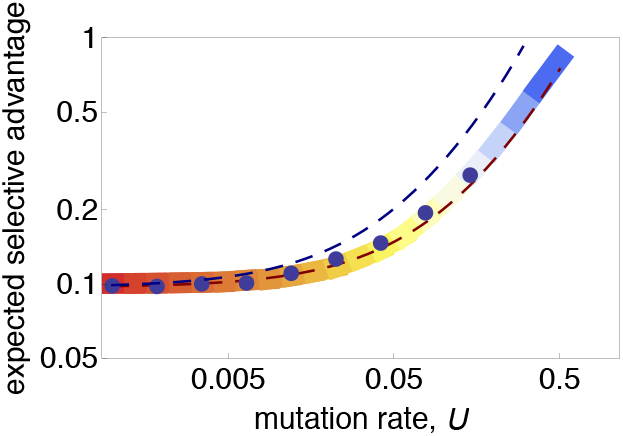
Expected selective advantage of surviving beneficial mutations as a function of deleterious mutation rate. Survival probabilities are computed in the same way as in Figure 4, employing Equation (5) and fixing *s_b_* = *s_b_*. Survival probability at each *s_b_* was multiplied by 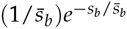, the probability that a mutation of selective advantage *s_b_* is produced. This product resulted in a unimodal curve, which we then normalized to create a probability density; from this, the expected selective advantage was computed. The thick solid curve has a color gradient indicating log probability of occurrence of beneficial mutations with the plotted selective advantage 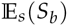 (red high, blue low). Expected selective advantages are approximated (dashed lines) as the expected value of a Gamma distribution with shape parameter 2 (Rozen *et al*. 2002; Gerrish and Lenski 1998) and truncated from below at *e^U^* − 1; this expected value is 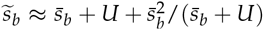. Parameters are: *s_b_* = 0.05, and *sd* = 0.03. Employing the same parameters, each blue dot plots the average selective advantage of surviving beneficial mutations from 15,000 simulations. We note the absence of blue dots above *U* ≈ 0.3, owing to the fact that none of the 15,000 beneficial mutations survived above this mutation rate (alluding to the “fixation threshold” derived below).

### Mutation rate that maximizes production of surviving beneficial mutations

The recruitment rate of beneficial mutations increases with genomic mutation rate, but because of lineage contamination, the survival probability of beneficial mutations decreases with genomic mutation rate. Therefore, there must exist a genomic mutation rate that maximizes the rate of production of surviving beneficial mutations. Setting *∂_U_U* 〉*p_svl_*〈 = 0 and solving for *U*, we find this maximum production rate occurs at mutation rate 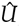, bounded as:

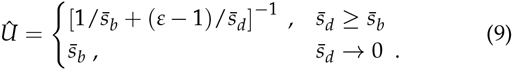

Figure 7 compares the foregoing predictions to simulation results and shows them to be quite accurate. The smallest value of 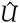 is achieved when 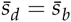, resulting in the bounds:

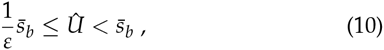

again displaying a curious independence of 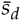. For reasoning similar to that given after Equation (8), the upper bound on 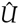 is approximated under the wider range of circumstances.

**Figure 7.**
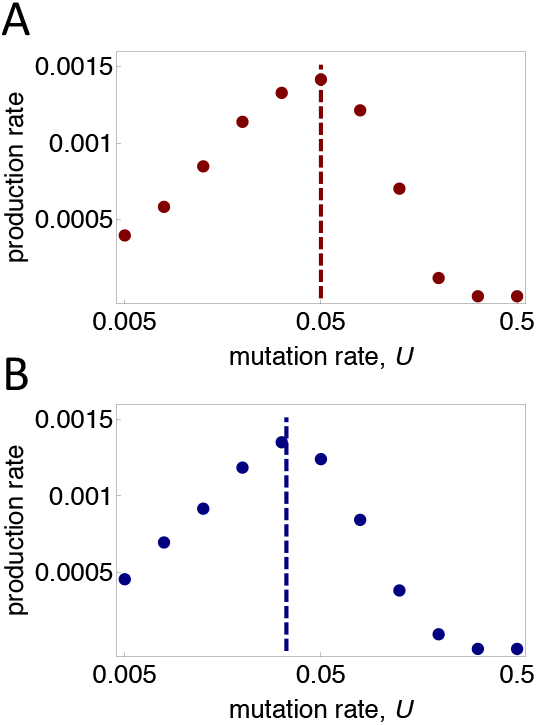
Theory accurately predicts mutation rate 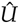 that maximizes production rate of surviving beneficial mutations (9). Beneficial mutations have mean selective advantage 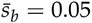. Each point represents 15,000 stochastic, individual-based simulations. (A) The weak-deleterious case 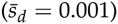 for which 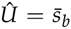 (vertical dashed line). (B) The strong-deleterious case 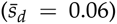 for which 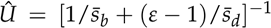 (vertical dashed line).

## Effects of excessive mutation

The lineage contamination effect we describe will increase with increasing deleterious mutation rate. When the mutation rate is high enough, this effect can cause the within-population mutational meltdown of many newly-arising beneficial lineages causing a reduction in competition and clonal interference. At even higher mutation rates, this effect can suppress most or all newly-arising beneficial lineages, resulting in the partial or complete cessation of adaptive evolution.

### Clonal interference threshold

Evolutionary dynamics may be naturally partitioned into different regimes, depending on the recruitment rate of beneficial mutations. At very low rates of recruitment of beneficial mutations, adaptive evolution proceeds through isolated selective sweeps – a regime that has been dubbed the “periodic selection” regime (Sniegowski and Gerrish 2010). As the recruitment rate of beneficial mutations increases, a point is reached at which two or more alternative beneficial mutations may coexist and compete for fixation (the “clonal interference” regime; Gerrish and Lenski (1998)). As recruitment rate of beneficials continues to increase, it may become likely that competition occurs not among single beneficial mutations but among genotypes carrying multiple beneficial mutations (the “multiple-mutations clonal interference” regime; Desai and Fisher (2007); Desai *et al*. (2007)).

What much of this previous work failed to account for (c.f. Orr (2000); Bachtrog and Gordo (2004b)) was the fact that, as beneficial recruitment rate increases via an increase in overall genomic mutation rate, the rate of deleterious mutation increases in parallel. The findings we have presented so far suggest an intriguing implication of this parallel increase: whereas beneficial recruitment rate increases linearly with genomic mutation rate, survival probability of beneficial mutations decreases exponentially with genomic mutation rate. This fact suggests that, at high genomic mutation rates, the effects of lineage contamination can overwhelm the increased production of beneficial mutations, such that the effective recruitment rate of beneficials (i.e., the rate of production of *surviving* beneficial mutations) can decrease as mutation rate increases further. As mutation rate increases, therefore, adaptive evolution may eventually revert to a regime in which it proceeds only through isolated selective sweeps; put differently, the population may revert from one of the clonal interference regimes back to the periodic selection regime at high mutation rates.

As delineated in Sniegowski and Gerrish (2010), the clonal interference regime is entered when a second, alternative beneficial mutation is likely to be produced on the ancestral background before the first, or focal, beneficial mutation becomes fixed. Mathematically, this transition occurs when *Nμ*〈*p_svl_*〉 ln(*Ns_b_*/2) / *s_b_* = 1 (SI), where *μ* = *cU* is the beneficial mutation rate and *c* is thus the ratio of beneficial to deleterious rates. In previous work, the transition considered was that which occurs as very low mutation rates increase, and 〈*p_svl_*〉 was taken to be some function (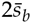 or some variant thereof) that was independent of *U*. Here, we have shown that, at high mutation rates, 〈*p_svl_*〉 can depend strongly on U. We define the “clonal interference threshold” to be the critical mutation rate above which adaptive evolution *reverts* from a clonal interference regime back to the periodic selection regime. This threshold is defined as:

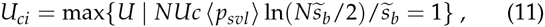

where 〈*p_svl_*〉 is defined by (8), *c* is the ratio of numbers of potential beneficial to deleterious mutations, i.e., *c* = *μ*/*U*, and 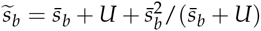, the expected selective advantage of beneficial mutations that survive (SI).

### Fixation threshold

The critical selective advantage below which a beneficial mutation does not survive increases approximately linearly with mutation rate when lineage contamination is considered in isolation, and faster than linearly when background selection is also accounted for. In contrast, the fittest mutation produced by a population has a selective advantage that increases approximately linearly with the *log* of the mutation rate. This necessarily implies that, as mutation rate increases, eventually a point will be reached at which even the selective advantage of the fittest beneficial mutation will not be sufficient to overcome the effects of lineage contamination. This point defines the “fixation threshold”, and its existence follows from the fact that the critical selective advantage required and the maximum selective advantage produced by a population have qualitatively different relationships with mutation rate.

The fixation threshold is exceeded when no beneficial mutation produced by a population has a selective advantage strong enough to survive the effects of lineage contamination. Concretely, in a given interval of time *τ*, we suppose a population produces a total of *n* beneficial mutations; then, the fixation threshold is defined as the mutation rate that ensures extinction of even the fittest of these mutations. This critical mutation rate, which defines the fixation threshold and which we will denote by *U_f_*, thus ensures the extinction of all *n* beneficial mutations produced with specified probability *p_c_*; it is given by:

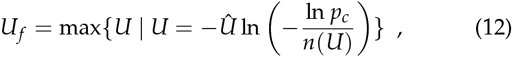

where *Û* is given by (9) and bounded by (10), *n*(*U*) = *NUcKŝ_b_τ*, and *τ* denotes the relevant time period; for example, to compute the mutation rate at which, over a time period of 5000 generations, all fixations will be suppressed with probability 95%, we set *τ* = 5000 and *p_c_* = 0.95.

**Figure 8.**
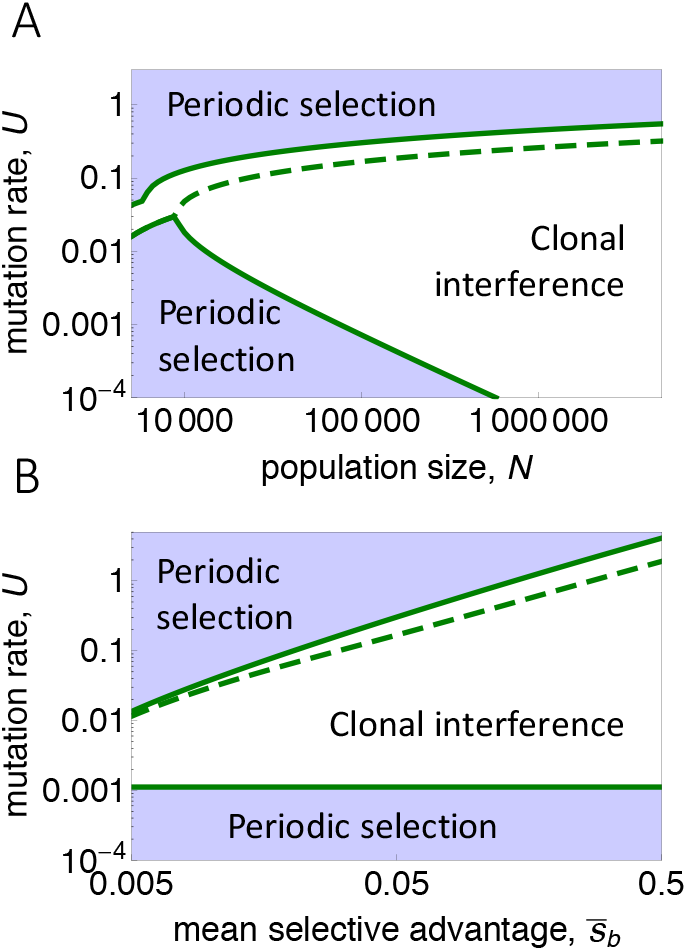
Clonal interference threshold. Mutation rates *U_ci_* that define boundaries between clonal interference and periodic selection regimes: 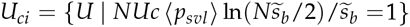. The greater of the two solutions (upper green solid curves, given by Equation (11)), represents the transition, as high mutation rate increases, from a clonal interference regime back to a periodic selection regime (the *clonal interference threshold*); the lesser of the two solutions (lower green solid curves) represents the first transition, as low mutation rate increases, from a periodic selection regime into the clonal interference regime. The dashed green curve plots the lower bound (for the case 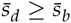) on the threshold. (A) Parameters are: 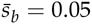 (upper solid curve); 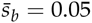 and 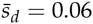 (lower dashed curve). (B) Parameters are: *N* = 500,000 (upper solid curve); *N* = 500,000 and 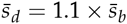 (lower dashed curve).

**Figure 9.**
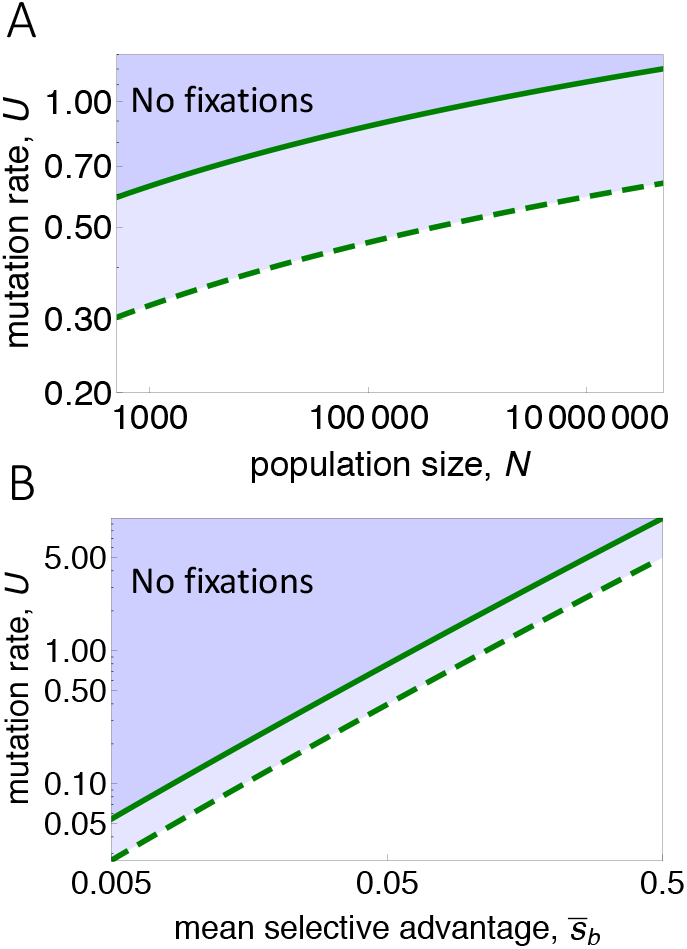
Fixation threshold. Green solid and dashed curves plot upper and lower bounds of *U_f_*, as given by (12). (A) Parameters are: 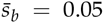 (upper solid curve); 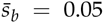 and 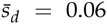 (lower dashed curve). (B) Parameters are: *N* = 500, 000 (upper solid curve); *N* = 500, 000 and 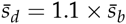 (lower dashed curve).

**Figure 10.**
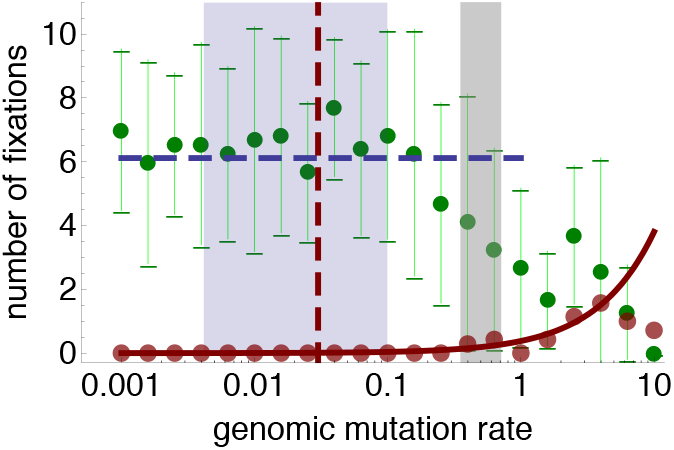
Total numbers of fixations in simulations of population size 50000 over the course of 2000 generations, as a function of genomic mutation rate (green dots), with error bars of 1.96 standard deviations. Deleterious and beneficial mutation rates were fixed fractions 0.5 and 0.0005 of the genomic mutation rate, respectively. Blue horizontal dashed line indicates the number of fixations above which is the clonal interference regime; light-blue shaded area delineates the mutation rates for which the population is predicted to be in the clonal interference regime. Red vertical dashed line plots the upper-bound mutation rate at which the production rate of surviving beneficial mutations is maximized (closely corresponding the maximum number of fixations), given by Equation (9). Gray vertical bar plots the mutation-rate interval bounded by the predicted “fixation threshold” bounds (12), predicting suppression of all fixations with probability *p_c_* = 0.95. Red solid curve plots the theoretical neutral expectation: numbers of neutral fixations expected to occur in the same time period; red points plot numbers of neutral fixations in separate simulations. Above the predicted “fixation threshold” (gray bar), numbers of fixations are not statistically distinguishable from the neutral expectation (green bars overlap with red curve).

## Discussion

### Summary

Evolutionary interactions between linked deleterious and beneficial mutations have received increasing attention in recent years. It is now well accepted, for example, that background selection caused by the continual rain of deleterious mutations into regions of low recombination decreases the fixation probability of beneficial mutations (Charlesworth *et al*. 1993; Peck 1994) and decreases nucleotide diversity (Stephan 2010; Kim and Stephan 2000; Birky 1988; Keightley and Otto 2006); moreover, theoretical and empirical studies have shown that selective sweeps of beneficial mutations can cause the fixation of linked deleterious mutations (McDonald *et al*. 2011; Bachtrog and Gordo 2004b; Good and Desai 2014; Hartfield *et al*. 2010; C W Birky and Walsh 1988). To date, work in this area has been focused on populations with relatively low genomic mutation rates. In such populations, the key consideration in analyzing the interaction between beneficial and deleterious mutations is the number of deleterious mutations *already* present in the linked genomic background on which a new beneficial mutation arises. In the current paper, we have focused, in contrast, on populations in which genomic mutation rates may be very high: We have examined the possibility that the genomic background on which a beneficial mutation arises can become progressively contaminated with newly arising deleterious mutations even as the beneficial mutation spreads into the larger population. Our work is motivated in part by numerous studies indicating that adapting asexual populations tend to evolve high mutation rates through genetic hitchhiking (Sniegowski *et al*. 1997, 2000; Johnson 1999a; Elena and Sanjuán 2005; Gentile *et al*. 2011; Söderberg and Berg 2011; M’Gonigle *et al*. 2009; Raynes *et al*. 2011) and by the substantial literature that has been devoted to the question of when the genomic mutation rate will be sufficiently high to cause population extinction (Gerrish *et al*. 2007, 2013; Gerrish and Sniegowski 2012; Bull and Wilke 2008; Bull *et al*. 2007; Springman *et al*. 2009; Biebricher and Eigen 2005; Eigen 2002, 2000, 1971; Eigen and Schuster 1977).

### Multiple beneficial mutations

Our multitype branching process model assumes that beneficial mutations occur infrequently enough that acquiring a second beneficial mutation in linkage with the focal beneficial mutation is improbable in the time required for the focal mutation to either survive or go extinct. In reality, it might be the case that multiple beneficial mutations arise on the same background and sweep to fixation, collectively overcoming the lineage contamination effect.

To assess the strength of our assumption, we studied the effects of allowing additional beneficial mutations to arise at different rates within the lineage founded by the focal beneficial mutation. To this end we varied the parameter *c*, introduced above and defined as: *c* = *μ*/*U*, or the ratio of numbers of potential beneficial to deleterious mutations. We assessed the effects of doing so in both an extension of our analytical model and in simulations.

To assess the effects of additional within-lineage beneficial mutations on lineage contamination in isolation, we extended our multitype branching process model so as to allow a beneficial lineage to acquire a second beneficial mutation with the same selective advantage *s_b_* as the first. This additional beneficial mutation could be acquired during reproduction with probability 1 – *e*^−*cU*^. Figure S7 compares, for different values of *c*, survival probabilities of a single beneficial lineage in an otherwise homogeneous population (lineage contamination only) as a function of the deleterious mutation rate. It is apparent from this figure that, for reasonable values of *c*, there is minimal quantitative difference in survival probabilities and only a slight increase in the apparent threshold. This indicates that our assumption of no additional within-lineage beneficial mutations is a weak assumption. Mathematically, there is a qualitative difference in that, for *c* > 0, the critical deleterious mutation rate above which a beneficial lineage becomes extinct almost surely (the “hard” threshold) is twice what it is without the additional beneficial mutation (i.e., 2 ln (1 + *s_b_*)). Practically, this is of little consequence, however, because the survival probabilities are typically minuscule for mutation rates in the region between the hard threshold for which no additional beneficial mutation is allowed (ln (1 + *s_b_*)) and the hard threshold for which one additional beneficial mutation is allowed (2 ln (1 + *s_b_*)). And, while not shown here, survival probabilities become even smaller in regions between higher thresholds that allow more beneficial mutations.

To assess the effects of additional within-lineage beneficial mutations arising in evolving populations, where both background selection and lineage contamination are operating, we performed simulations in which there was technically no limit on the number of additional beneficial mutations. Figure 5 plots survival probabilities computed from simulations, for the cases *c* = 0, 0.001, and 0.01, and would seem to indicate, again, that our original assumption of no additional within-lineage beneficial mutation is a very weak assumption. Survival probabilities such as those plotted in Fig. 5 were computed for a range of different parameters and in all cases, for what we considered to be reasonable beneficial-to-deleterious ratios (*c* ≤ 0.01), survival probabilities were essentially unaffected by the incorporation of additional beneficial mutations.

### Independence of our results from the selective effects of deleterious mutations

The critical mutation rate above which lineage contamination in isolation ensures extinction of a beneficial mutation, derived in the first section (1), depends only on the selective advantage of the focal beneficial mutation; it does not depend on the selective disadvantages of deleterious mutations. In the next section, when we incorporate background selection, most of the solution bounds we derive are also independent of the selective disadvantages of deleterious mutations. These results stand in contrast to some previously published results that have focused primarily on the effects of background selection. Particularly striking is the contrast between our results, which find a surprising lack of dependence on 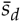, and the results of Orr (2000), which instead find a surprising lack of dependence on 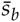. For example, we find that 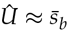, whereas Orr finds that 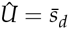. A thorough exploration of the relevant parameter space and assessment of Orr’s result is found in Johnson and Barton (2002b). The single factor that accounts for the qualitative discrepancy between our results and Orr’s is lineage contamination: when only background selection is accounted for, 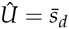, yet when lineage contamination is also accounted for, 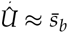. These “opposite” results shine a light on the impact of lineage contamination generally.

### Lineage contamination in nature

The effects of lineage contamination only become significant under linkage and relatively high mutation rates. While we have focused exclusively on the case of asexuality, lineage contamination should also operate in organisms that undergo some form of genetic exchange as well: the fitness of a newly-arising beneficial mutation will be eroded at a faster rate than the same linkage region in the rest of the population – on principles similar to those studied here. The requirement of high mutation rates would seem to restrict the relevance of our findings to organisms like RNA viruses, although the evolution of high mutation rates has been predicted (Gerrish *et al*. 2007) and increasingly reported in natural (Matic *et al*. 1997) and laboratory (Sniegowski *et al*. 1997; Shaver *et al*. 2002; Denver *et al*. 2009; Wichman 2005; Pal *et al*. 2007; Gentile *et al*. 2011; Chao and Cox 2008; Cox and Gibson 1974) populations of RNA and clonal DNA organisms and in somatic (esp. cancerous) cells. Indeed, the process we have analyzed has been implied in conjunction with background selection as a mechanism that can slow the evolution of tumors (McFarland *et al*. 2013; Solé 2004).

### Lineage contamination, mutational meltdown, and lethal mutagenesis

As alluded to in the presentation of our branching process model, lineage contamination may be thought of as within-population mutational meltdown. If this meltdown is induced by treatment of a population with a mutagenic agent, then it may be thought of as within-population lethal mutagenesis. Put differently, our findings may be thought of as the population-genetic analogues of these processes. Indeed, one of the processes we model – Muller’s ratchet in a growing beneficial lineage – is similar to previous models of Muller’s ratchet in freely-growing populations (Fontanari *et al*. 2003; Bull *et al*. 2007; Bull and Wilke 2008). Our work differs from these previous studies, however, in that we model the fitness erosion of a growing lineage within the context of a larger population.

In a recent experiment, lethal mutagenesis failed to cause extinction in a laboratory population of the bacteriophage T7 (Springman *et al*. 2009) because the accumulation of deleterious mutations opened up new genetic pathways that could increase fitness, i.e., it increased the number of available beneficial mutations. The theory we present here may offer insight into what would be required to thwart the evolutionary rescue afforded by these newly-available beneficial mutations. In particular, our “fixation threshold” might offer an appropriate quantitative guideline for the mutation rate required.

### Lineage contamination and the error threshold

There is an intriguing relationship between our findings and predictions of “error threshold” models (Eigen 2002, 1971; Eigen and Schuster 1977; Biebricher and Eigen 2005; Bagnoli and Bezzi 1998; Nowak and Schuster 1989; Bonhoeffer and Stadler 1993). Generally and somewhat loosely speaking, an error threshold is a critical mutation rate, *U_e_t*, above which all genotypes deterministically converge to the same *equilibrium* frequency, independent of their fitness (in the absence of mutational biases).

#### Single-peak model

The simplest model of the error threshold – the so-called “single peak” model – assumes that there is a single fittest genotype of fitness 1 + *s_b_* (the beneficial mutant) and all other (mutationally accessible) genotypes have fitness equal to one (Wiehe 1997; Tejero *et al*. 2011), i.e., the fitness landscape has a set of two possible fitness classes {1,1 + *s_b_*}. This fitness landscape is obviously unrealistic; its original conjecture may have been based on the fact that many other such “phase transition” phenomena are robust to severe model simplification. When the number of possible genotypes may be assumed to be infinite, the error threshold is *U_et_* = ln(1 + *s_b_*) (Wiehe 1997). Curiously, while the assumed fitness landscapes are very different, this critical mutation rate is identical to the one we derive for lineage contamination in isolation (1).

#### Multiplicative model

The set of possible fitness classes on the “multiplicative” fitness landscape is {(1 + *s_b_*)(1 – *s_d_*)^*i*^ ∀ *i* ∈ [0, *D*]}, where *D* is the maximum number of deleterious mutations allowed. Curiously, for the case *D* < ∞ and *s_b_* = *s_d_*, the error threshold is *U_et_* = *s_b_* (we note that for small *s_b_, U_et_* ≈ ln(1 + *s_b_*), in agreement with the “single-peak” model), whereas when *D* = ∞, there is no error threshold: *U_et_* = ∞ (Wiehe 1997). Oddly, our lineage contamination model corresponds most closely to the case *D* = ∞, for which there is no error threshold, but there is a lineage contamination threshold.

### Concluding remarks

Wittingly or not, the presence of lineage contamination has been implicit in many previous models of mutation-induced fitness erosion. To our knowledge, however, it has not previously been modeled in isolation, as a process separate from background selection. Our theoretical framework partitions these two processes, and allows lineage contamination to be scrutinized separately from other processes. We find, for example, that newly-arising beneficial mutations can be driven extinct almost surely by lineage contamination whereas background selection alone cannot ensure their extinction.

## Acknowledgements

We thank Thomas Bataillon, Guillaume Martin, Alan Perelson, Nick Hengartner, and Thomas Burr for helpful discussions. SP received financial support from the “Soutien à la recherche des jeunes maîtres de conférences” program at the Université Paris-Est Créteil. PG carried out much of this work in, and received financial support from, visiting faculty programs at Aarhus University, Denmark, and at University of Montpellier (Labex/CeMEB/ISEM), France. PS and PG received financial support from NASA grant NNA15BB04A.

## Supporting Information

### Beneficial lineages in an initially homogeneous population: multitype branching process model

Our stochastic model is a discrete-time multi-type branching process with infinite set of types 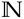, where a type 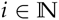 corresponds to the number of acquired deleterious mutations. Since we do not assume back mutations, the branching process is irreducible. We recall that the vector 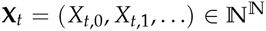 describes the composition of the population at time 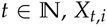, *X_t,i_* being the number of individuals in the population at time *t* carrying *i* deleterious mutations. This branching process can be entirely described by its offspring generating function **f** = (*f*_0_, *f*_1_, …), defined for each 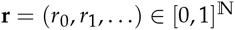 by 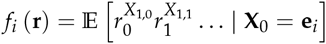, where **e**_*i*_ denotes the vector 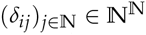 and *δ* stands for the Kronecker delta. By construction, for each 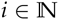 and each 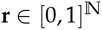, *f_i_* only depends on *r_i_*, *r*_*i*+1_, … and

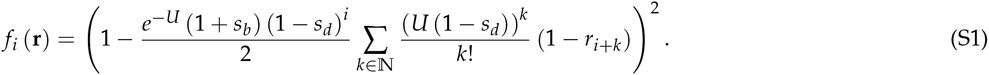

#### Mean demographic dynamics of each sub-population

The mean matrix 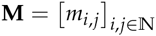 of the branching process, defined such that 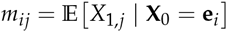 (mean number of individuals of type *j* produced by one individual of type *i*) is upper triangular with, for each *i*, 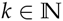,

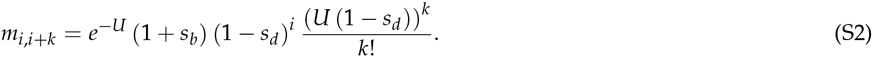

Moreover, for each 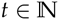, the mean matrix of the process at time *t* corresponds to the *t*-th power of **M**. We denote 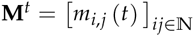. It satisfies 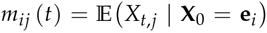 (mean number of individuals of type *j* at time *t* produced by one individual of type *i*). The matrix is upper triangular and is such that for each *i*, 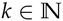,

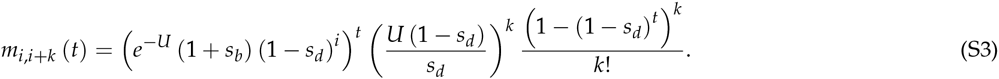

From this we deduce that the mean size of each sub-population at time 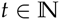 is given by

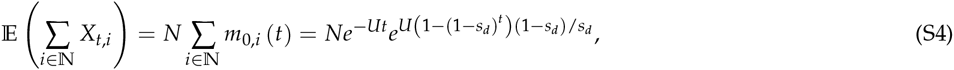

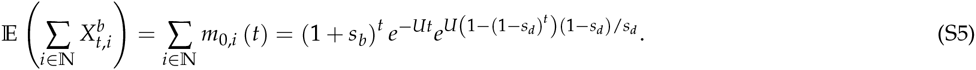

**Figure S1.**
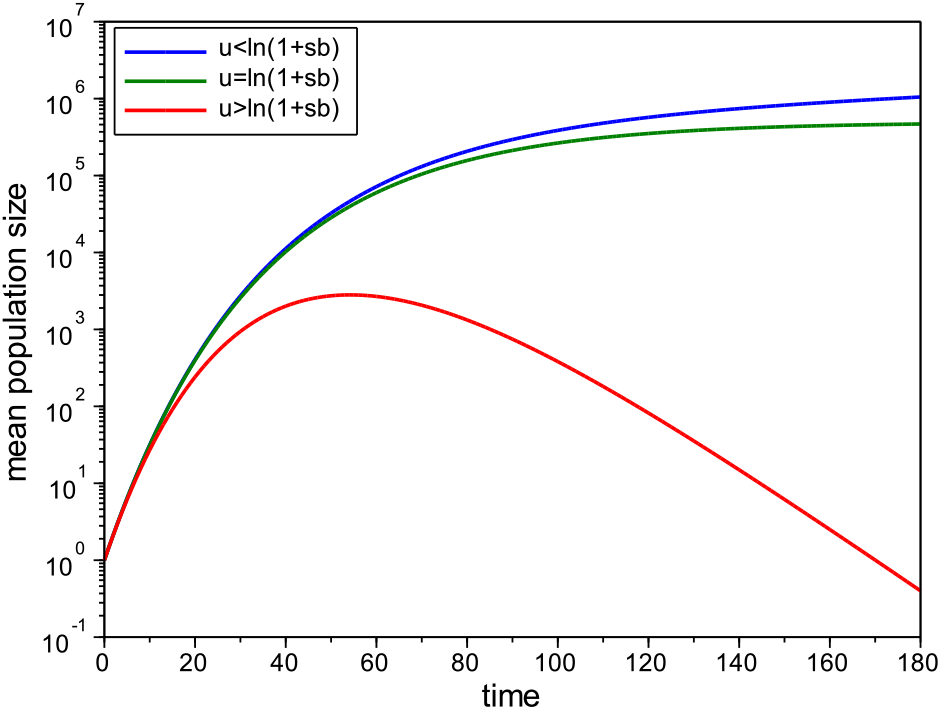
Evolution over time of the mean population size stemming from one single beneficial lineage, with *s_d_* = 0.03, *s_b_* = 0.5, and *U* = 0.04 (resp. *U* = ln(1 + 0.5), *U* = 0.5). We use a large value for *s_b_* to show that a beneficial lineage can grow to a large size before being driven extinct by lineage contamination (red curve).

**Figure S2.**
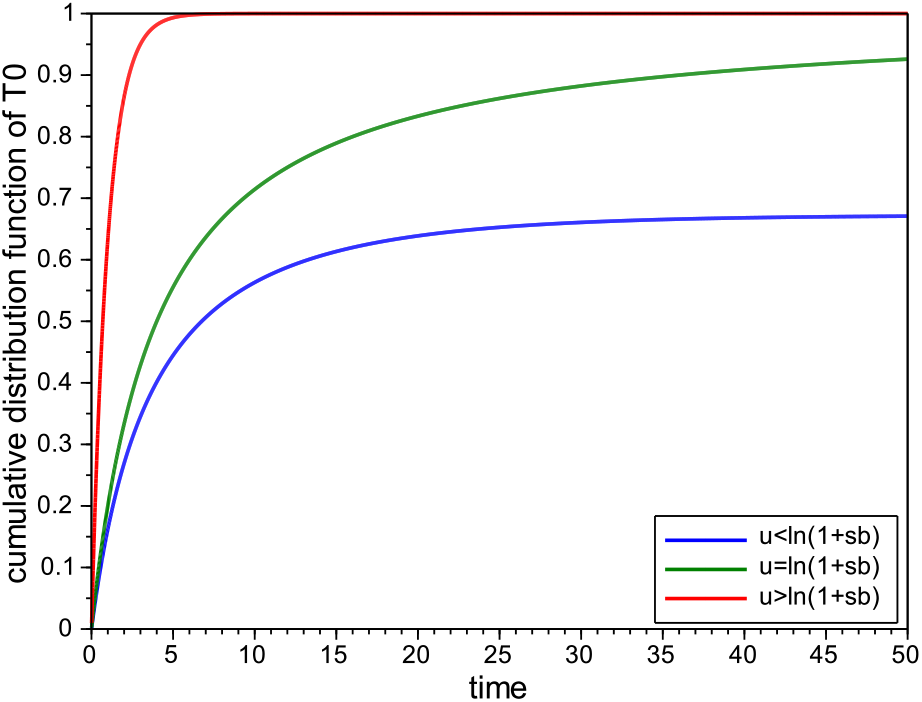
Cumulative distribution function of the first least-loaded class extinction time *T*_0_, for one single beneficial lineage, with *s_b_* = 0.1, *s_d_* = 0.03, and *U* = 0.001. (resp. *U* = ln(1 + 0.1), *U* = 10).

**Figure S3.**
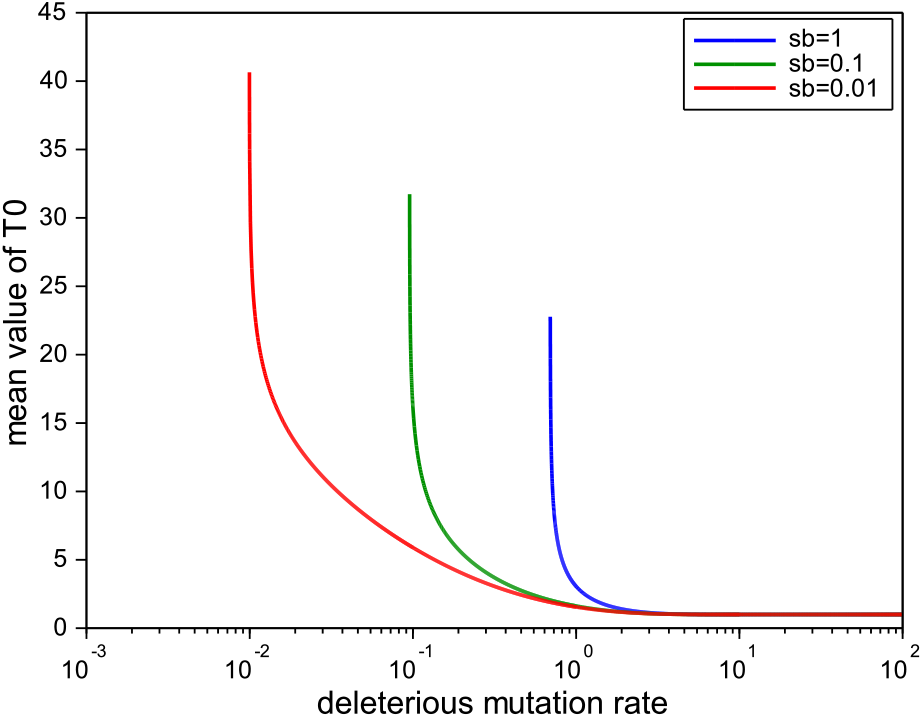
Mean value of the first least-loaded class extinction time *T*_0_, for one single beneficial lineage, with *s_d_* = 0.03. For *U* ⩽ ln(1 + *s_b_*), this mean value is infinit. ()

**Figure S4.**
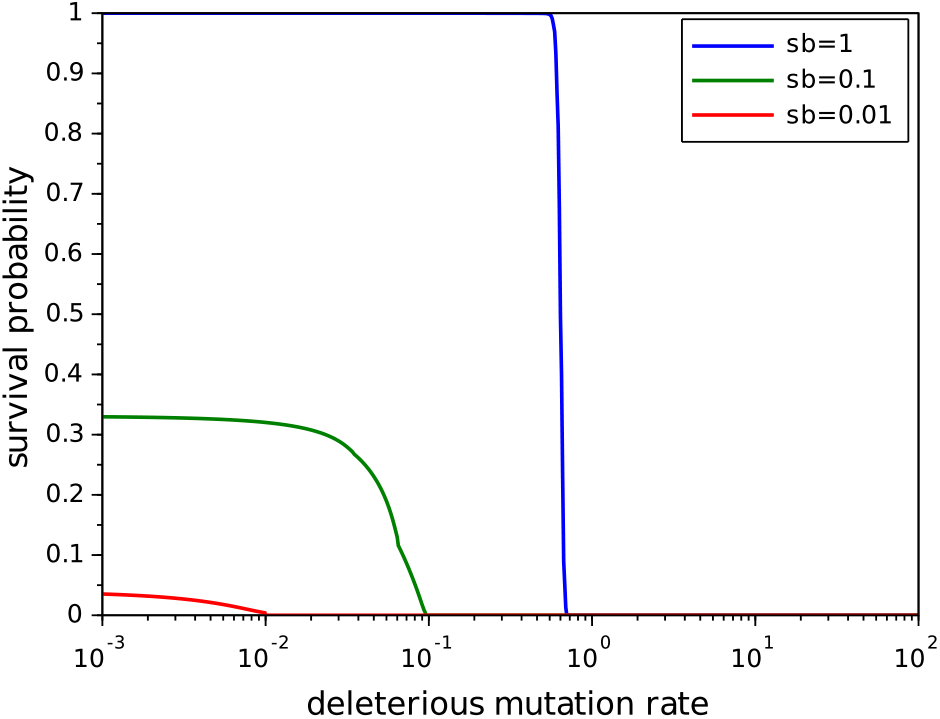
Survival probability 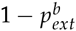 of a single beneficial lineage, with *s_d_* = 0.03, as a function of the deleterious mutation rate.

**Figure S5.**
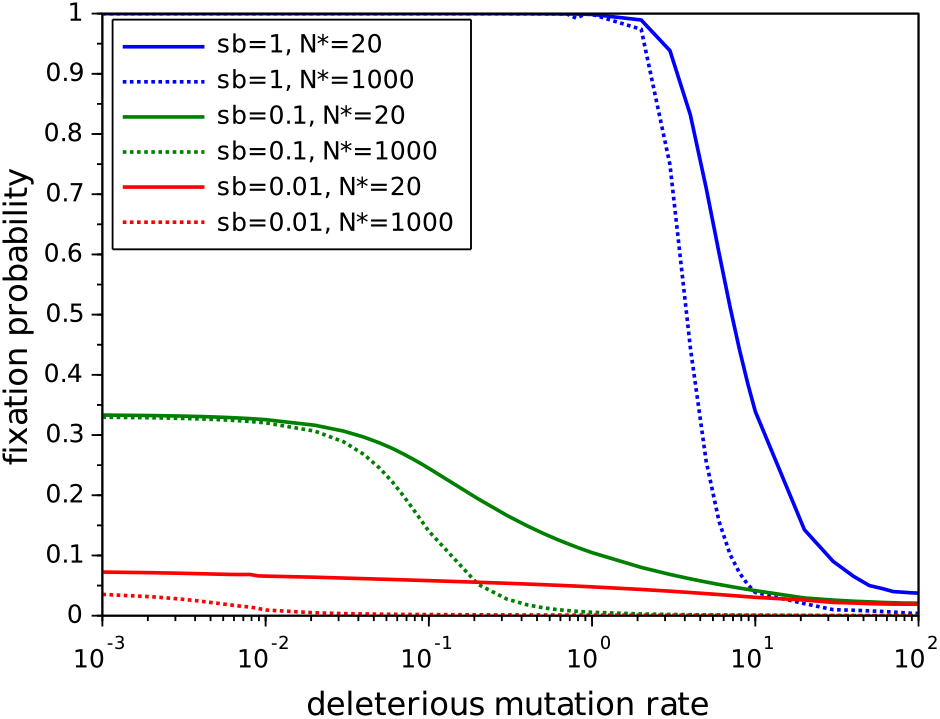
Fixation probability *p_fix_* of a single beneficial lineage in a wild-type population of initial size *N*, with *s_d_* = 0.03. Here *N*\* = *N* + 1 corresponds to the initial size of the total population.

**Figure S6.**
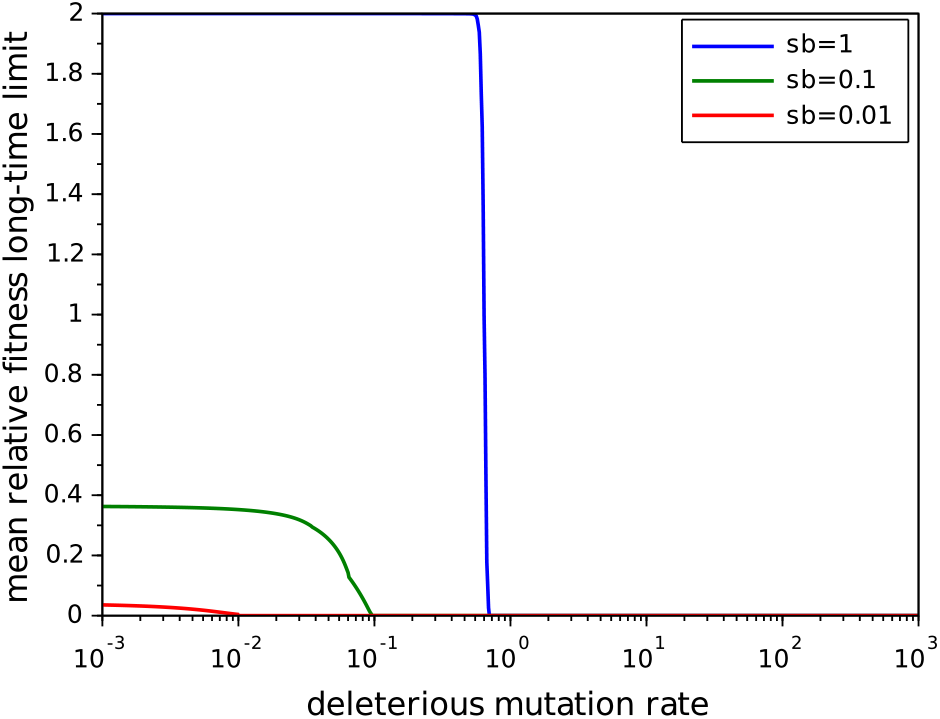
Long-term limit of the mean relative fitness of a single beneficial lineage in a large wild-type population, with *s_d_* = 0.03.

### Extinction probabilities

**Proposition 1**.

- *For any U* ⩾ 0 *and* 0 < *s_b_*, *s_d_* < 1, *the two following extinction probabilities of a single beneficial lineage* 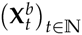 *are equal*:

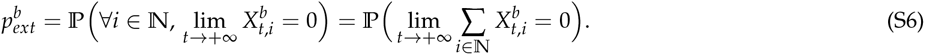

*Moreover*,

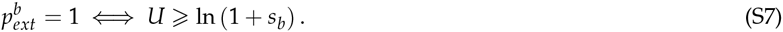
- *For any U* ⩾ 0, 0 < *s_d_* < 1 and 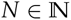, *a neutral population* 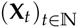 *of initial size N becomes almost surely (a.s.) extinct*:

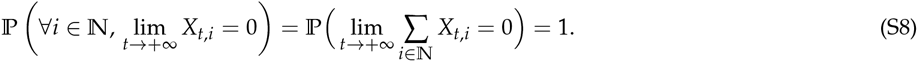

#### Proof.

The following proof holds for any initial population size and any *s_b_* ⩾ 0. In what follows the notation 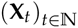 consequently stands indifferently for a neutral population of initial size *N* ⩾ 1 with *s_b_* = 0, or for a single beneficial lineage with *s_b_* > 0. If not mentioned otherwise, the probabilities in this proof are computed conditionally on the event {**X**_0_ = *N***e**_0_}.

The extinction of the whole population implies the extinction of every type, hence the obvious inclusion 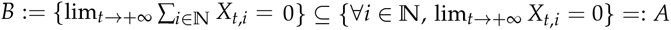. However, the simultaneous survival of the whole population and extinction of all the types imply the existence of at least one infinite line of descent with one descendant whose type is indefinitely increasing. We thus obtain by the Markov property that

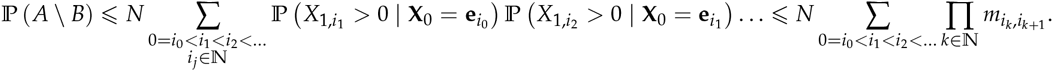

Since for each strictly increasing sequence 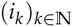 with *i*_0_ = 0 we have *i_k_* ⩾ *k*, it comes by (S2) that

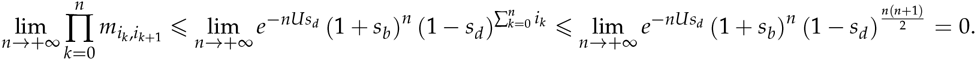

Hence 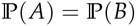, which in turn implies (S6) and the first equality in (S8).

Moreover, 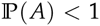 if and only if there exists some 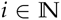 such that 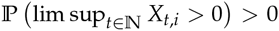. For each 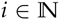, let us denote 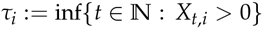. By the Markov property and thanks to the independence of the initial lineages we can write

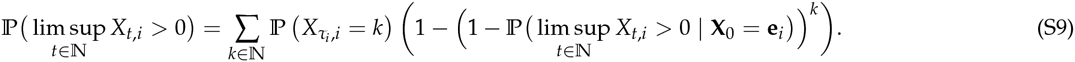

Since a mutant of type 0 can produce one mutant of type 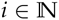 with positive probability, one can find for each *k*, 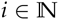 an event of the form {**X**_0_ = *N***e**_0_, …, **X**_*t*−1_ = *k***e**_0_, **X**_*t*_ = *k*_e__*i*_} ⊆ {X_*τ_i_,i*_ = *k*} occurring with positive probability, hence implying that 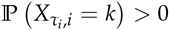. Consequently, (S9) implies that 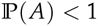 if and only if there exists some 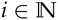 such that 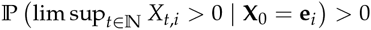. Seeing the infinite-type branching process 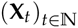 as a branching random walk on 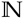 as in Zucca (2011), the last assertion means that there exists 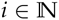 such that so-called local survival of the branching walk at *i* occurs with positive probability, starting with one individual at *i*. By Theorem 4.1 in Zucca (2011), local survival at *i* occurs with positive probability if and only if 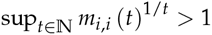, which by (S3) is equivalent to *e^−U^* (1 + *s_b_*) (1 − *s_d_*)^i^ > 1. Hence 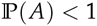 if and only if *e^−U^* (1 + *s_b_*) > 1, which proves (S7) and the second equality in (S8).

#### Numerical computation of the extinction probability

We use here the fact that the extinction probability *p_ext_* can be obtained as the limit of the extinction probability of a multi-type branching process with a finite number of types, when the number of types tends to infinity. We show moreover that this limit value is actually reached for a finite number of types, which consequently leads to an exact computation (see (S11)) rather than an numerical approximation.

First note that if *U* ⩾ ln (1 + *s_b_*), then *p_ext_* = 1 by Proposition 1. We thus assume from now on that *U* < ln (1 + *s_b_*). For each 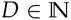, we shall associate with 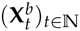 the process 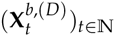 in which none of the individuals with more than *D* acquired deleterious mutations are counted. Then 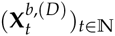 is a multi-type branching process with finite set of types {0, …, *D*} and with offspring generating function **f**^(*D*)^ given for each **r** ∈ [0,1]^*D*+1^ and *i* ∈ {0, …, *D*} by

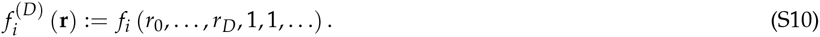

Let us define its extinction probability vector 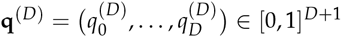 where for each *i* ∈ {0, …, *D*},

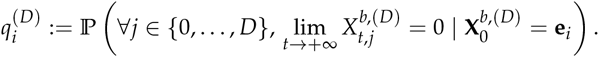

Let us moreover define 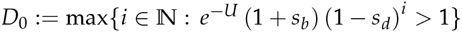. We then have

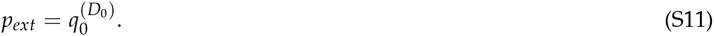

Indeed, we know from Hautphenne *et aL*. (2013) that **q**^(*D*)^ converges as *D* → +∞ to the “partial” extinction probability of 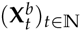, which by (S6) is equal to its “global” extinction probability. Hence in particular 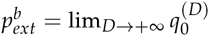. Moreover, since for each *i* > *D*_0_ we have *m_ii_* < 1, a process starting from one mutant of type *i* almost surely becomes extinct, which implies that for each *D* ⩾ *D*_0_, 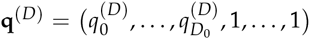. Moreover, it is known (see e.g. Satz V.1.4 in Sewastjanow (1975)) that for each 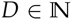, **q**^(*D*)^ is the smallest non-negative fixed point of **f**^(*D*)^. From what precedes this implies that for each 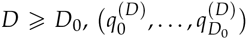 is the smallest non-negative solution of the system

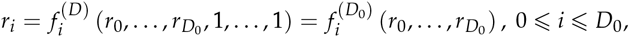

which is known to be **q**^(*D*_0_)^. Consequently, for each *D* ⩾ *D*_0_, 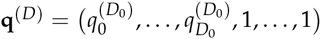, leading to the desired result (S11).

### Fixation probabilities

Similarly as for the extinction probability we approximate *p_fix_* by the fixation probability of one single beneficial lineage in a neutral population of initial size *N*, when none of the individuals carrying more than *D* deleterious mutations are counted. For this purpose we consider for each 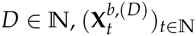 and 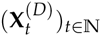 the multi-type branching processes with offspring generating function (S10) (with *s_b_* = 0 for the process 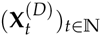), and such that 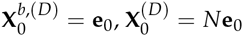. Let 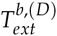 and 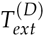 their respective extinction times. Then, according to Hautphenne *et aL*. (2013), for 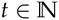 fixed and *D* → +∞, 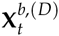 (resp. 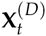) almost surely monotonically converges to 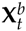 (resp. **X**_*t*_). Hence 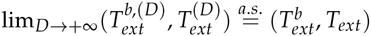 and in particular

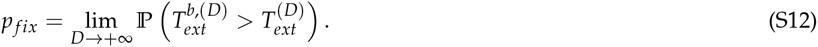

Let us describe how to compute the probability 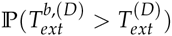 for any fixed 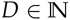. First, it can be shown by standard arguments that the multi-type branching process with finite number of types 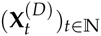 almost surely becomes extinct, i.e. 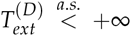. Consequently,

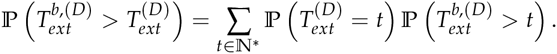

Note for instance that for each 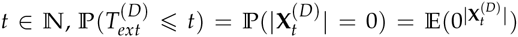, and that by independence of the *N* lineages, 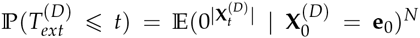. Furthermore (see e.g. Athreya and Ney (2004) Section V.1), the generating function 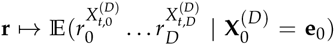 of the branching process at time 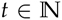 is given by the *t*-th iterate of **f**^(*D*)^ defined by (S10), with *s_b_* = 0. We denote **f**^(*D*),0^ = *id* and for each 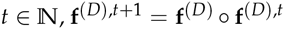. Therefore, 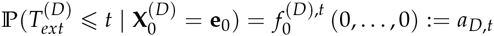, where *s_b_* = 0 in (S1). Similarly, taking *s_b_* > 0 in (S10), 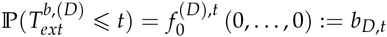. Hence we finally obtain that

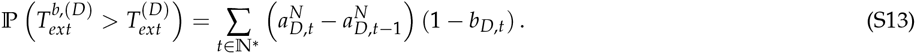

### Fitness dynamics of the beneficial lineages within a population

#### Large population approximation

By construction, the population 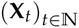 can be written as the sum of *N* independent and identically distributed copies of a branching process with offspring generating function (S1) (with *s_b_* = 0), and initial state **e**_0_, which we write 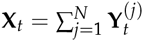. The strong law of large numbers combined with (S3) then implies that for each 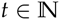, *t* < *T_ext_*,

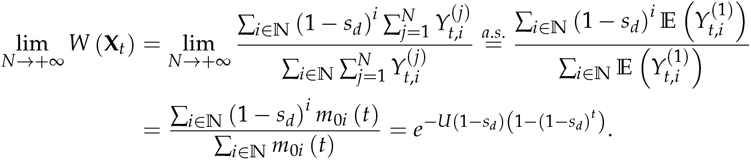

The same limit would be obtained for 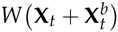, hence the approximation

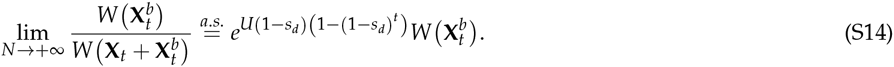

#### Long-time limit of the mean relative-fitness

Let us first assume that *U* ⩾ ln (1 + *s_b_*). By Proposition 1, 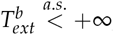, which approximating the relative fitness by its almost sure limit (S14) leads to

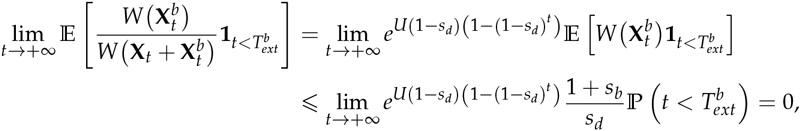

Since 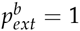, we have proven the desired result

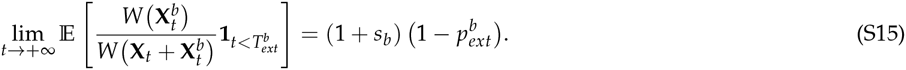

Let us now assume that *U* < ln (1 + *s_b_*). In this part of the proof we approximate 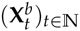 by the multi-type branching process 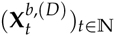 with finite set of types {0, …, *D*}, 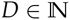, introduced previously (see (S10)). We assume 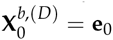. Note that its mean matrix corresponds to the truncated matrix [*m_i,j_*]_0⩽*i,j*⩽*D*_, with largest eigenvalue *m*_00_ > 1. Hence 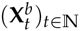 is not irreducible, and we thus make use of Theorem 2.1 in Kesten and Stigum (1967) for reducible branching processes with largest eigenvalue greater than 1 to deduce the existence of some one-dimensional random variable *Z*^(*D*)^ with 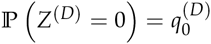 such that for each *i* ∈ {0, …, *D*},

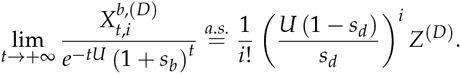

Defining the absolute fitness 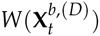 and the extinction time 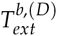 similarly as before we thus obtain that

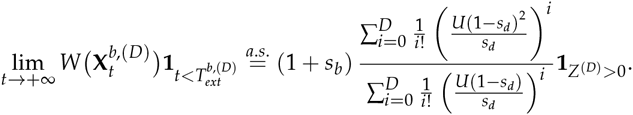

By the dominated convergence theorem, the previous convergence also holds in *L*^1^. Consequently, using the fact that 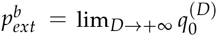,

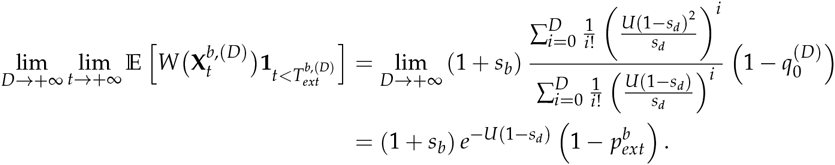

Using again the approximation (S14), we finally obtain

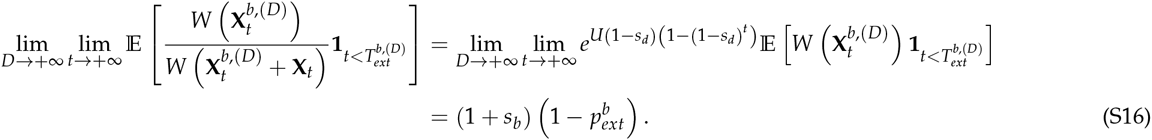

#### Upper and lower bounds of the mean relative fitness

Our goal is to provide upper and lower bounds of the mean relative fitness for each 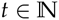, which as detailed below are both numerically computable by iteration of suitable generating functions. Defining 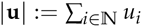 for any 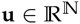 we have for each 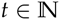, using (S14),

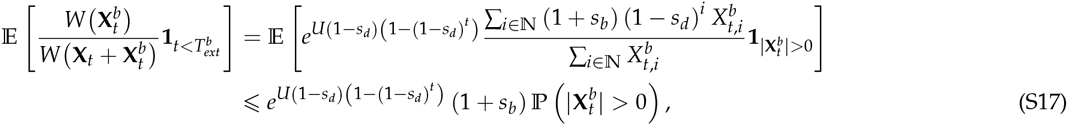

and, by Cauchy-Schwarz inequality,

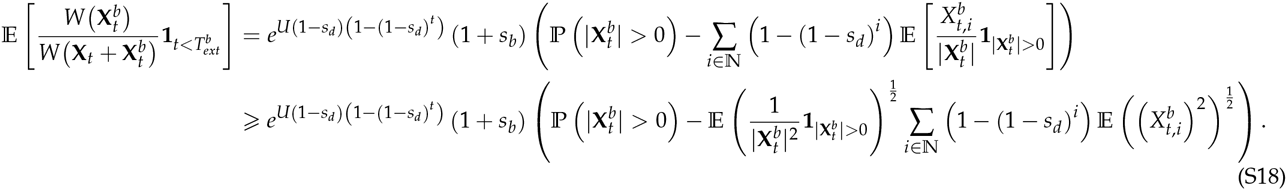

#### Numerical approximation of the mean relative fitness bounds

Similarly as before, let us for each 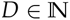 consider the branching process 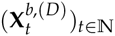 with offspring generating function **f**^(*D*)^ defined by (S10) (where *s_b_* > 0), and let us denote by **f**^(*D*),t^ its *t*-th iterate. Then the almost sure monotonous non-decreasing convergence of 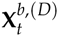 to 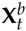, for 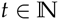 fixed, entails that the terms appearing in (S17)–(S18) can be for each 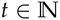 obtained as follows. First,

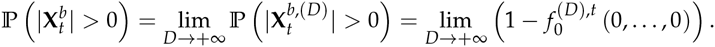

Second,

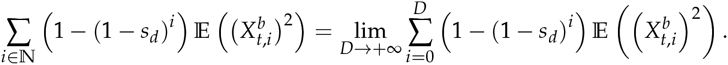

By construction, for each 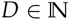 and each 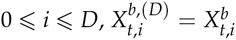. Hence, denoting for each *r* ∈ [0,1]

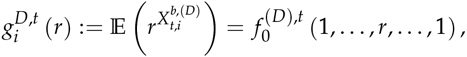

we obtain that

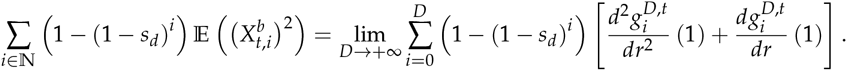

Finally, the dominated convergence theorem implies that

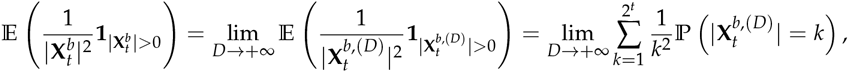

where for each 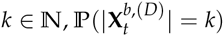 is the coefficient of *r^k^* in the polynomial function

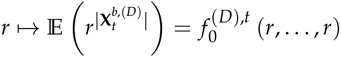

of degree 2^*t*^. This coefficient can be for instance obtained with a software such as Scilab via the tools pol and `coef`.

### Mutational meltdown of a beneficial lineage

We recall that in this section we model the random accumulation of deleterious mutations via the continuous-time analog 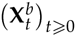 of the branching process 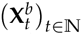 used until now. In this setting, each individual has a random lifespan which is exponentially distributed with parameter 1 (instead of a deterministic lifespan of one time-unit), and produces offspring at the end of the lifespan as previously. This modification allows us to benefit from the fact that the generating functions of continuous-time branching processes are solutions of differential equations, which will in our model lead to a computation of the cumulative distribution function of the fittest class extinction time. The following proposition provides the sequence 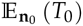, 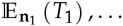 of the mean extinction times of the least loaded-classes, where 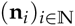 reflects the mean evolution of 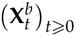, that is to say **n**_0_ = (1,0,0,…), and 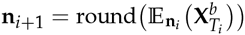, for each 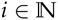. For each 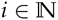, we define 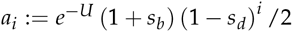.

**Proposition 2.**

- If *U* < ln(1 + *s_b_*),

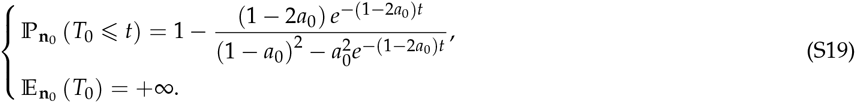
- If *U* = ln(1 + *s_b_*),

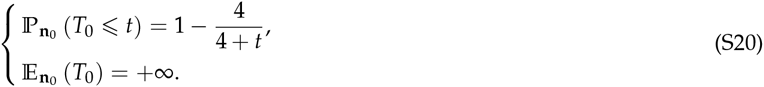
- If *U* > ln(1 + *s_b_*),

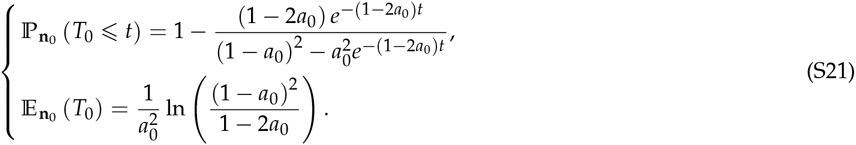

*More generally, for each* 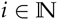 *and j* > i,

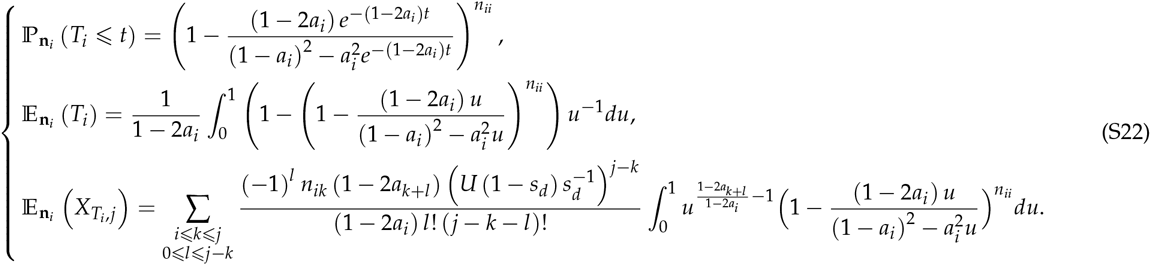

Note that this prove in particular that 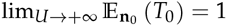, as we can see in Figure S3.

*Proof*. Let 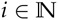 and 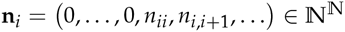. Note that under 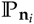, the law of the extinction time *T_i_* does not depend on *n_ij_, j* > *i*, since none of the individuals of type *j* can produce individuals of type *i*. Under 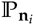, the extinction time *T_i_* actually has the same distribution as the extinction time of a single-type branching process 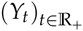 in which only the individuals of type *i* are counted, conditionally on {ϒ_0_ = *n_ii_*}. By construction, the offspring generating function of 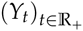 is given for each *r* ∈ [0,1] by *g*(*r*) = *f_i_* (*r*, 1,1,…) = (1 − *a_i_* + *a_i_r*)^2^. For each 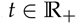, let us define 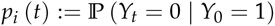. Then for each 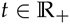,

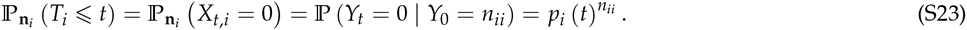

Moreover (see for instance III.3. in Athreya and Ney (2004)), 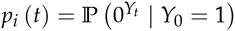 is solution of the differential equation

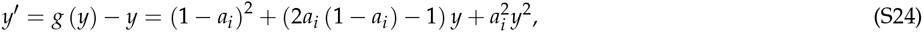

with the boundary condition *p_i_* (0) = 0. Note that (S24) is a Riccati equation for which one solution (*y*(*t*) = 1) is known.

If 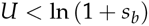 then 2*a*_0_ > 1 and solving (S24) leads to

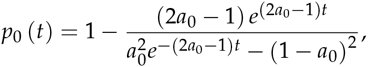

which combined with (S23) entails (S19). In particular, 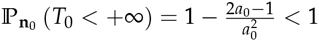 and thus 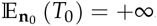.

If 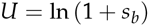 then 2*a*_0_ = 1 and (S24) leads to

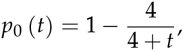

leading to (S20) and immediately implying that 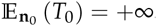.

Finally, if 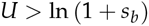 then for each 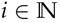, 2*a_i_* < 1, and solving (S24) leads to

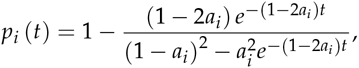

hence the first equality in (S21) and (S22). Since *T_i_* is a positive random variable, its first moment is given by 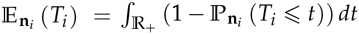. Integrating by substitution we obtain the second equality in (S21) and (S22). Note that a more explicit but rather bulky formula could be found for 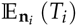, but for the sake of simplicity we choose to keep the definite integral form which is directly computable with a software such as Scilab. We moreover deduce from the first equality in (S22) that for each *j* > *i*,

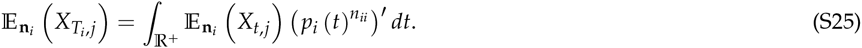

In the continuous-time setting, the mean composition 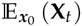 of the multi-type branching process at time 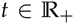, conditionally on {**X**_0_ = *x*_0_}, 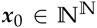, is given by the vector ***x***_0_*e*^(**M**−**I**)*t*^ (see for instance Athreya and Ney (2004)), where *e* denotes the matrix exponential. Hence by (S3) we obtain that for each 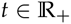,

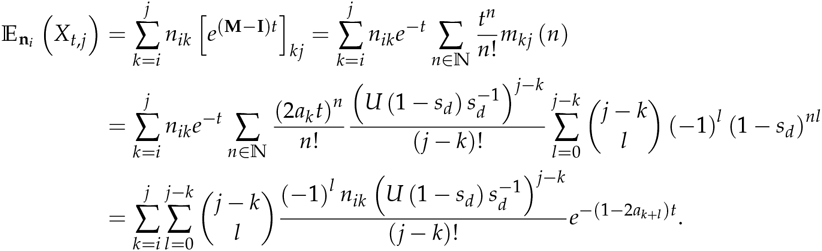

Integrating by part (S25) next leads to

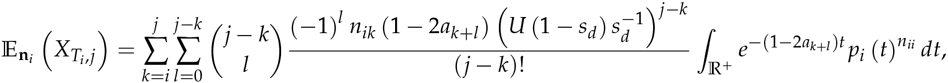

Finally, an integration by substitution entails the third equality in (S22), which is a definite integral easily computable with any suitable software.

### Beneficial lineages in an evolving population

#### Incorporating background selection

In the absence of beneficial mutations, an idealized (Wright-Fisher) population will over time reach a state in which individuals within the population carry a Poisson-distributed number of deleterious mutations with mean 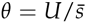, where *U* is deleterious mutation rate and 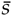 is geometric mean effect of deleterious mutations (Orr 2000; Haigh 1978b; Johnson 1999b). This now classical result breaks down, however, at sufficiently high mutation rate, *U*. Gessler (1995) points out that Poisson classes with frequency less than 1/*N*, where *N* is population size, (i.e., fitness classes in the tails of the Poisson distribution) will not exist: their frequency will be zero. While this observation is of little consequence where the right tail of the distribution is concerned, it is nevertheless quite consequential to the evolution of the population where the left (high fitness) tail of the distribution is concerned.

In fact the fittest *k* classes may disappear deterministically (left tail may be truncated) if their expected frequency is less than 1/*N*. Intuitively, this number may be computed as:

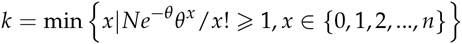

As the *k* fittest classes disappear due to mutation pressure, the evolving distribution of numbers of deleterious mutations tends toward a shifted Poisson:

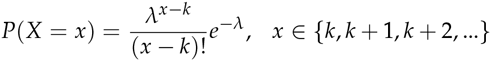

where *λ* = *θ* − *k*. Notably, this distribution has mean *θ* and variance *θ* − *k*, meaning that it is “under-dispersed” relative to the Poisson distribution: *V*(*X*) < *E*(*X*), where *X* is a random variable denoting number of deleterious mutations.

Gessler’s key insight was the perhaps counterintuitive subtlety that there is no guarantee, from the above derivations, that the zero class of the shifted Poisson distribution exists, i.e., that *e*^−*λ*^ ⩾ 1/*N*. The condition used to find *k* is that it be the smallest class in the Poisson distribution with parameter *θ* whose expected frequency exceeds 1/*N*. This condition, however, does not guarantee that this same class exists (has frequency greater than 1/N) in the new shifted-Poisson distribution with parameter λ = *θ* − *k*. Put differently, the fittest class in the new shifted distribution is not necessarily the class that carries *k* deleterious mutations.

Instead, to find the fittest, or ‘best’, class in the new distribution, we must find the smallest class whose expected frequency is greater than 1/*N* in a new truncated Poisson:

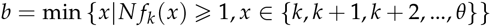

where *f_k_*(*x*) is a shifted Poisson truncated on the left at *k*:

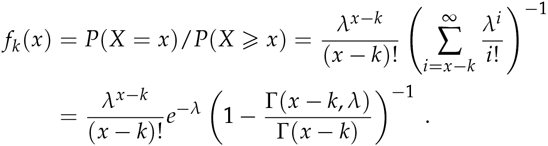

(The final expression is our own simplification of Gessler.)

#### Survival probabilities

In direct analogue to (2), we can immediately write down the survival probability of a beneficial mutation arising in a heterogeneous population. If the beneficial mutation in question has selective *s*_b_ and it arises on a genetic background carrying *j* deleterious mutations, then the initial growth rate of the beneficial lineage formed by this beneficial mutation will be:

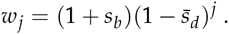

The survival probability of this beneficial mutation is bounded by:

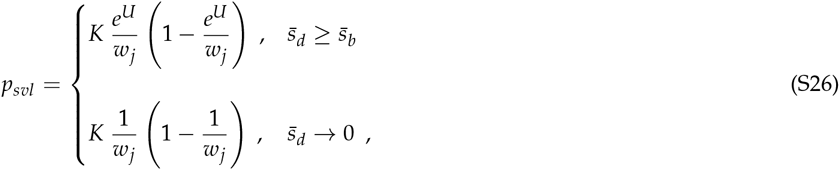

If we do not know the selective advantage of the beneficial mutation in question, and we do not know how many deleterious mutations are in the background upon which the beneficial mutation arises, we can employ the distributions of these quantities to compute ensemble survival probabilities of beneficial mutations as:

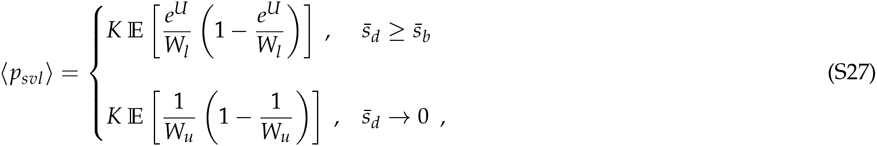

where *K* = 4 for binary fission, and *K* ≈ 2 for Poisson-distributed offspring; 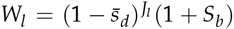 and 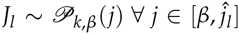; 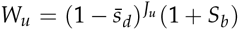 and 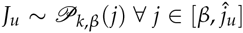; *S_b_* is also a random variable and, for each value of 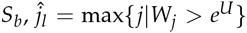 and 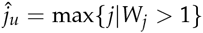, and *W_j_* is defined as 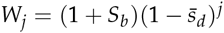.

##### Monte Carlo integration

Monte Carlo integration of (S27) was achieved by first drawing *S_b_* from an exponential distribution with mean 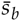, then computing 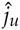 and 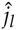, and finally computing a value for *p_sυl_*, and repeating. The average of the values computed for *p_sυl_* in this way was then taken to be the ensemble average 〈*p_sυl_*〉.

##### Distribution of beneficial mutational effects

We will rely on statistical arguments put forth by Gillespie (1991), after Smid and Stam (1975), that selection coefficients of beneficial mutations are exponentially distributed. While this assumption, and the statistical arguments upon which it is based, have been questioned and in some cases rejected based on different data analyses (Rokyta *et al*. 2008), we employ it here because of its far-reaching basin of attraction, for its simplicity, and because it is likely to hold approximately in the short term even if it does not ultimately hold exactly in the long term.

##### Refining the Haldane survival probability

Haldane derived what he called the “fixation probability” of a beneficial mutation of selective advantage *s_b_* to be approximately 2*s_b_*. In our terminology, this quantity is more accurately called the “survival probability”, as it does not take into account finite population size. Because we will be comparing our results to very large numbers of simulations, we would like to refine Haldane’s result a bit.

###### General expression

Haldane’s derivation starts with a result from single-type branching process theory, namely, that extinction probability of a Galton-Watson process is the smallest positive *x* that satisfies *x* = *h*(*x*), where *h*(*x*) denotes the probability generating function associated with the distribution of numbers of offspring produced by individuals in the population. To compute survival probability – the complement of extinction probability – we simply replace *x* with *y* = 1 − *x*. Survival probability is thus the largest *y* ∈ 2 [0,1] satisfying 1 − *y* = *h*(1 − *y*). Expansion yields

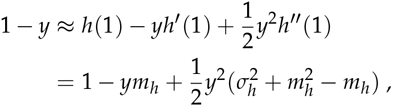

where *m_h_* = *h*′(1) is the mean number of offspring, and 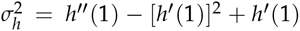 is the variance in number of offspring. Rearranging thus yields a general approximation for survival probability in the single-type case:

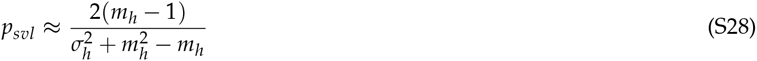

In this single-type case the mean number of offspring does not change over time and is given by *mh* = 1 + *s_b_*, resulting in:

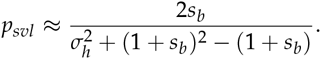

If it is the case that the quantity 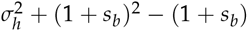 is close to one, then the survival probability may be approximated as *p_sυl_* ≈ 2*s_b_*, recovering Haldane’s classical result.

###### Poisson-distributed offspring

When numbers of offspring have a Poisson distribution, then we have 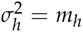, and (S28) becomes:

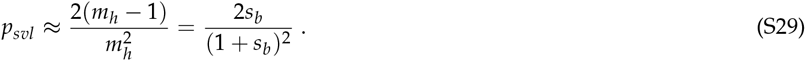

###### Bacteria (binary fission)

After reproduction and subsequent population culling, or sampling, the number of offspring a bacterium produces has a Binomial distribution with parameters *n* = 2 and *p* = *m_h_*/2, i.e., *h*(*x*) = [(1 − *m_h_*/2) + (*m_h_*/2)x]^2^. For this particular case, no approximation is necessary: 1 − *y* = [(1 − *m_h_*/2) + (*m_h_*/2)(1 − *y*)]^2^ may be solved exactly for *y*, yielding:

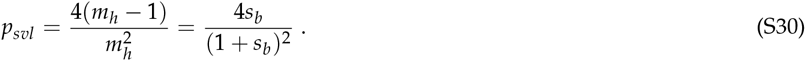

###### Expression used in main text

Results in the main text are presented in a way that can accommodate either of the foregoing two cases by implementing the following expression for single-type survival probability:

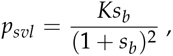

where *K* = 2 for Poisson-distributed offspring and *K* = 4 for bacteria (binary fission). When *s_b_* is replace with average selective advantage, 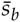, we introduce a new variable 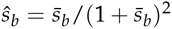, so that single-type survival probability may be written as:

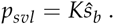

#### Approximation

##### Strategy and rationale

Rationale for our approximation strategy is based on the observation that, as *s_b_* increases, survival probability jumps from zero to something close to Haldane’s classical survival probability at *U* = ln *W*, where *W* is the relative fitness of the beneficial lineage; this is especially true for small 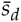.

Thus, we approximate survival probabilities as the product of the probabilities that, for a randomly chosen beneficial mutation: 1) the mutation has a selective advantage large enough to overcome the effects of lineage contamination, and 2) the mutation survives. For such beneficial mutations, the probability of 1) is the probability that *U* < ln *W*; and, the probability of 2) is our refined Haldane survival probability 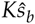.

##### Framework

To facilitate the following developments, we introduce a new function, 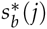, defined as follows. If a beneficial mutation arises on a background that contains *j* deleterious mutations, it has zero probability of survival unless its selective advantage exceeds a critical value, which is given by the function 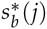, defined as:

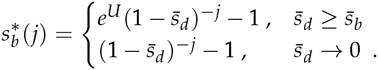

This function comes from the observation that zero probability of survival in (S26) is achieved when 1 − *e^U^*/*w_j_* ≤ 0 (i.e., when *e^U^* ≥ *w_j_*) for the case 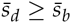, or when 1 − 1/*w_j_* ≤ 0 (i.e., when 1 ≥ *w_j_*) for the case 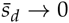. It is of interest, and will be used later on, to note that the ensemble average critical selective advantage is:

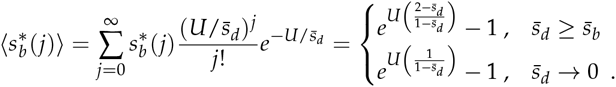

Based on the rationale outlined above for our approximation strategy, survival probability is thus:

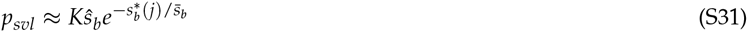

where 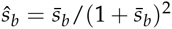, as derived above. Ensemble survival probability is thus:

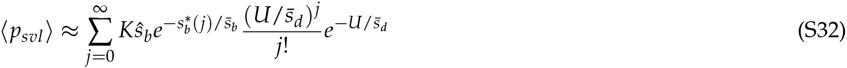

##### Lower hounds on ensemhLe survivaL probablity

To derive lower bounds on survival probability, we take the logarithm of (S31):

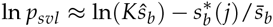

and write corresponding ensemble log survival probability:

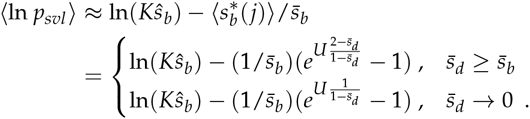

Jensen’s inequality provides:

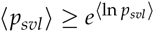

and thus yields exact minimums on both upper and lower bounds for the ensemble survival probability:

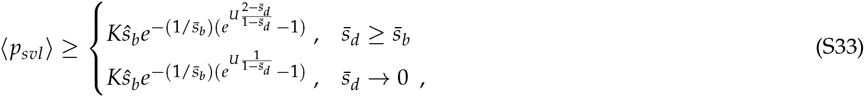

This expression is a bound-of-bounds and thus of questionable utility. Comparison with simulations, however, reveals the upper bound (for 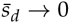) to be quite accurate; nevertheless, the lower bound (for 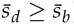) appears to be overly conservative.

##### Approximate ensemble survival probability

We now employ a different approach that avoids the use of Jensen’s inequality by making approximations that, at first blush, may appear somewhat crude; nevertheless, comparison with simulations shows this approach to work spectacularly well. We first note that typical values of 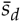 (even when larger than 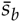) are likely to be small enough to permit the following approximations:

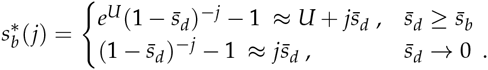

(S32) now becomes:

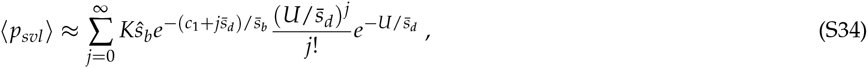

where *c*_1_ = *U* when 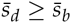, and *c*_1_ = 0 when 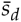 is small. This gives rise to approximate survival probabilities:

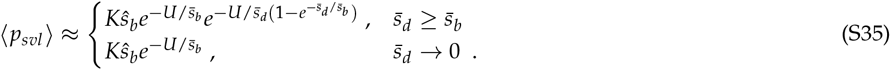

From here, it is apparent that the smallest value of 〈*p_svl_*〉 is achieved when 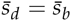, so that survival probability is bounded as:

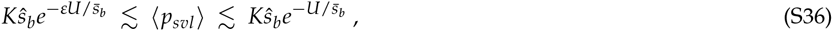

where *ε* = 2 – *e*^−1^ ≈ 1.63. Remarkably, the foregoing bounds on survival probability are independent of 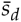.

#### Mutation rate that maximizes production of surviving beneficial mutations

We first note that the rate of production of surviving beneficial mutations will be proportional to *U*〈*p_svl_*〉. This is the only information we need. The genomic mutation rate that maximizes this rate of production is found by setting *•_U_U* 〈*p_sdl_*〉 = 0 and solving for *U*, we find this maximum production rate occurs at mutation rate *Û*, given by:

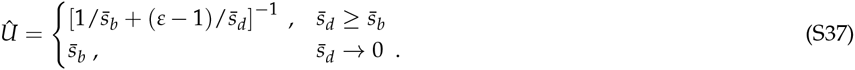

The smallest value of *Û* is achieved when 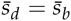, resulting in the bounds:

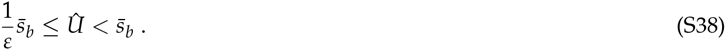

#### Clonal interference threshold

As delineated in Sniegowski and Gerrish (2010), the clonal interference regime is entered when a second, alternative beneficial mutation is likely to be produced on the ancestral background before the first, or focal, beneficial mutation becomes fixed. If the focal mutation has selective advantage *s_b_* then, given that it survives stochastic sampling in the first few generations (genetic drift), the dynamics of its frequency, *x*(*t*), is described by the differential equation,

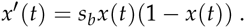

Conditioning on surviving drift (Barton 1995), the initial condition is:

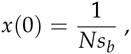

yielding solution:

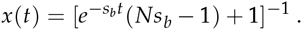

At this point, we refine our definition of “fixation” of the focal beneficial mutation as being the point at which half of the population carries the mutation. The rationale for this refinement is that beyond this point, when more than half of the population carry the mutation, it becomes more likely that the next beneficial mutation to arise will appear not on the wildtype background but on the background carrying the focal beneficial mutation. Fixation, defined in this way, will occur *τ* generations after the appearance of the focal mutations, where *x*(*τ*) = 1/2. Thus,

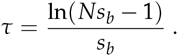

We are interested in knowing whether an alternative beneficial mutation will arise in the interval (0, *τ*) *on the wildtype background*. Letting *y*(*t*) denote wildtype frequency at time *t*, we have that

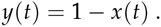

The expected number of alternative beneficial mutations produced in the wildtype subpopulation is:

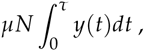

where *μ* = *cU* is beneficial mutation rate, and:

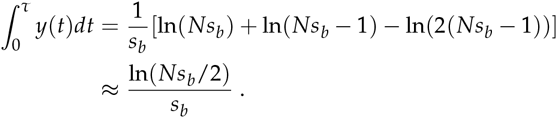

The expected number of alternative beneficial mutations *produced* on the wildtype background is thus:

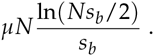

The expected number of alternative beneficial mutations produced, however, is different from the number of *competing* beneficial mutations, because many of those produced will not survive the effects of genetic drift and the accelerated accumulation of deleterious mutations (lineage contamination). Letting random variable *M* denote the number of *competing* beneficial mutations, then *M* has a Poisson distribution with expectation:

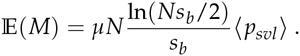

A population is in the clonal interference regime when at least one competing beneficial will arise on average; this criterion is thus met when:

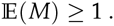

The only remaining issue is to determine the value of *s_b_* to be used in the foregoing expressions. We will use 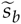 denote this value. To let 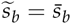 would be erroneous, because beneficial mutations that fix will typically have higher-that-average selective advantages. And to survive the effects of lineage contamination, we have established that the selective advantage must be at least *e^U^* – 1. In previous work (Rozen *et al*. 2002; Gerrish and Lenski 1998), we find that selective advantages of beneficial mutations that survive drift will be Gamma distributed with shape parameter 2. Fortunately, on the border between clonal interference and periodic selection regimes, we expect only a single competing beneficial mutation, and we therefore do not have to account for the effects of clonal interference on selective advantages of beneficial mutations that fix. In light of the foregoing observations, we have:

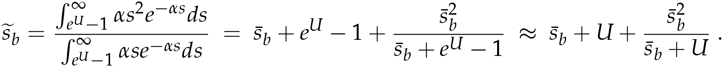

It is readily verified that, as *U* → 0, 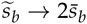, as expected.

Finally, we note that there will typically be two solutions to the equation 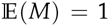, which defines the borders between clonal interference and periodic selection regimes. The smaller of these two solutions defines the border, as low mutation rate increases, from a periodic selection regime into a clonal interference regime. The larger of these two solutions defines the border, as already high mutation rate increase further, from a clonal interference regime back into a periodic selection regime. This second transition is the direct result of lineage contamination and is what we have called the “clonal interference threshold”.

#### Fixation threshold

The probability that a randomly chosen beneficial mutation does *not* survive is:

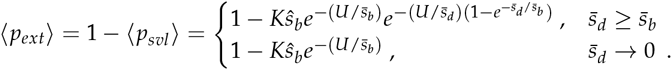

where *k* ∈ (1, *ε*]. The probability that *n* beneficial mutations do not survive is thus:

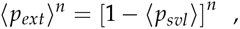

which may be approximated by:

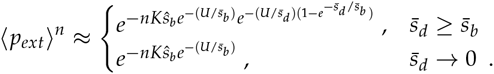

The fixation threshold is obtained by finding the maximum *U* that satisfies:

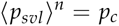

where *p_c_* is the probability with which one wishes to ensure that no fixations occur. The “fixation threshold” is exceeded when no beneficial mutation produced by a population has a selective advantage strong enough to survive the effects of lineage contamination. Concretely, in a given interval of time *τ*, we suppose a population produces a total of *n* beneficial mutations; then, the fixation threshold is defined as the mutation rate that ensures extinction of even the fittest of these mutations. This critical mutation rate, which defines the fixation threshold and which we will denote by *U_f_*, thus ensures the extinction of all *n* beneficial mutations produced with specified probability *p_c_*; it is given by:

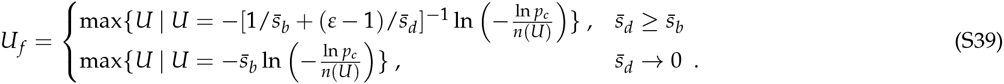

More compactly, this may be rewritten as:

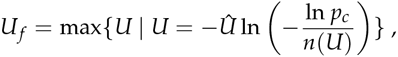

where *Û* is defined and derived above, and *n*(*U*) = *NUcKŝ_b_τ*, and *τ* denotes the relevant time period; for example, to compute the mutation rate at which, over a time period of 5000 generations, all fixations will be suppressed with probability 95%, we set *τ* = 5000 and *p_c_* = 0.95.

#### Multiple beneficial mutations

Our multitype branching process model assumes that beneficial mutations occur infrequently enough that acquiring a second beneficial mutation in linkage with the focal beneficial mutation is improbable in the time required for the focal mutation to either survive or go extinct. In reality, it might be the case that multiple beneficial mutations arise on the same background and sweep to fixation, collectively overcoming the lineage contamination effect. Figure S7 plots survival probabilities computed from an extension of our multitype branching process model that allows for the occurrence of a second beneficial mutation within the lineage formed by the first, or focal, beneficial mutation.

**Figure S7.**
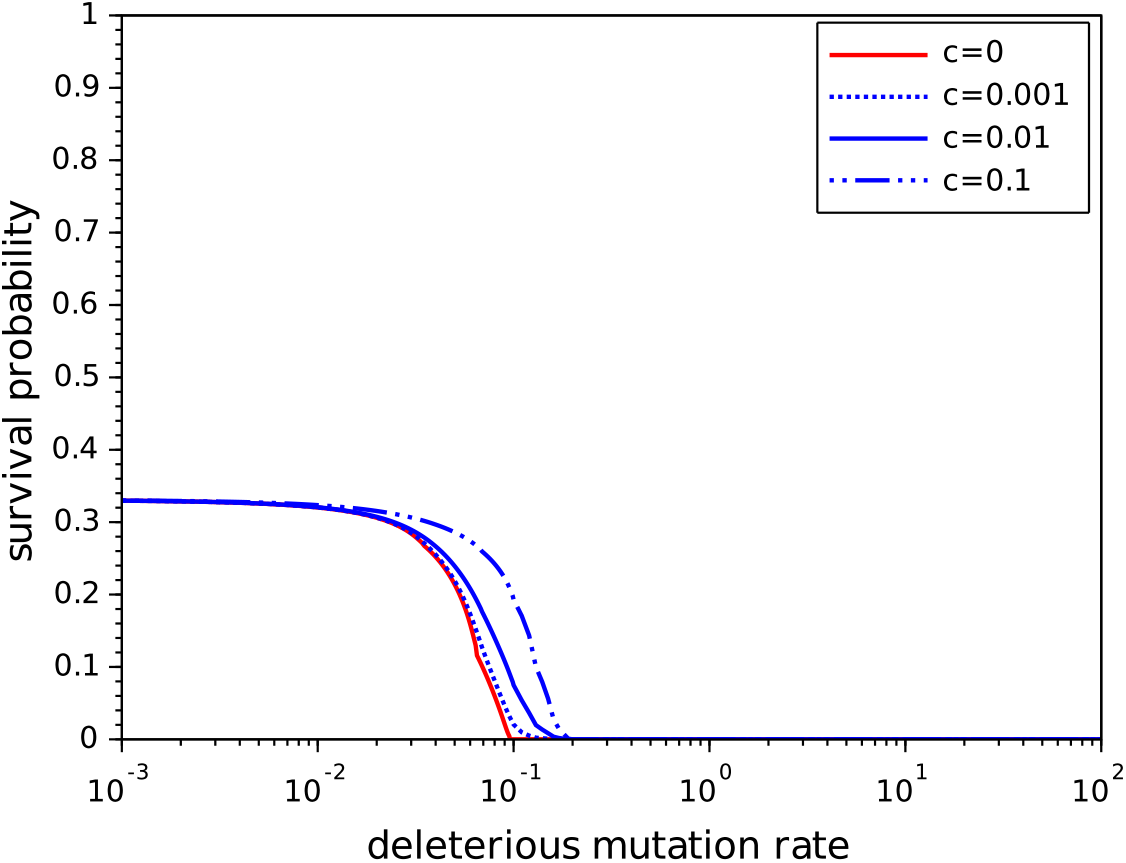
Survival probability of a single beneficial lineage, potentially acquiring a second beneficial mutation with probability 1 – *e*^−cU^. Here *s_b_* = 0.1 and *s_d_* = 0.03.

#### Simulations

Our simulations describe the evolution of a population whose size at time *t* is *N_t_*. At time *t*, the *i^th^* individual in the population has fitness *W_t,i_*, so simulations model the evolution of the vector **W**_*t*_ = (*W*_*t*,1_ *W*_*t*,2_,…, *W_t,N_t__*). To compute vector **W**_*t*+1_, the first step is replication: each individual is granted a number of offspring, *Y_t,i_*, proportional to that individual’s fitness: *Y_t,i_* is a Poisson random variable with mean 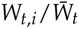, where 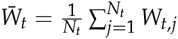. If individual *j* in the population at time *t* + 1 is an offspring of individual *i* in the parent population (at time *t*), then its fitness is computed by applying 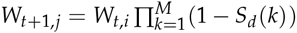, where *i* = 1,2,…, *N_t_*, each *S_d_*(*k*) is an exponentially-distributed random variable with mean *s_d_*, and *M* is a Poisson-distributed random variable with mean *U*.

##### Single beneficial lineage in an otherwise homogeneous population

To study the effects of lineage contamination in isolation, we simulate the growth of a beneficial lineage in an initially homogeneous population by imposing the initial conditions: *W*_0,1_ = (1 + *s_b_*) and *W*_0,*i*_ = 1 for *i* = 2,3,…, *N*_0_. These simulations closely approximate the process modeled by our branching process approach described above, but they allow us to study the effects of relaxing certain assumptions.

##### SingLe heneficiaL Lineage in a heterogeneous popuLation initiaUy at mutation-seLection balance

In other simulations, we analyze the growth of a single beneficial lineage occurring at random (on any background) in a population initially at mutation-selection balance. Here, initial conditions are: 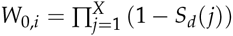, where *X* is a Poisson-distributed random variate with mean *U*/*s_d_*, and the *S_d_*(*j*) are exponential random variates with mean *s_d_*, and *W*_0,*K*_ = *W*_0,*K*_ (1 + *s_b_*) where *K* is an integer randomly chosen in the interval [1, *N*_0_].

##### MuLtipLe heneficiaL mutations in a heterogeneous popuLation

These simulations assume a deleterious genome of infinite length and a binary (bit-string) beneficial genome of length 96 (three binary “genes” each of length 32). Each of the 96 positions on the beneficial genome was assigned a fixed fitness effect of mutation at that position. Initially, this set of selection coefficients was created by drawing at random from an exponential distribution with mean 0.03. For purposes of consistent comparison, we used the same set of available beneficial mutations for all such simulations.

##### IsoLating Lineage contamination and hackground selection processes

In simulations, we isolated the effects of the lineage contamination and background selection processes by allowing one of these processes to operate while suppressing the other. To examine the effects of lineage contamination alone, we suppressed background selection by not allowing deleterious mutations to arise during the replication of an individual with zero beneficial mutations. To examine the effects of background selection alone, we suppressed lineage contamination by not allowing deleterious mutations to arise during the early growth of beneficial lineages. And of course, to examine the combined effects of lineage contamination and background selection, we allowed equally indiscriminate mutation (we suppressed neither process).

